# A Multi-omic Atlas of Human Choroid Plexus in Alzheimer’s Disease

**DOI:** 10.1101/2025.11.23.690014

**Authors:** Tristan J. Philippe, Denis R. Avey, Nicola A. Kearns, Himanshu Vyas, Sashini De Tissera, Jishu Xu, Bernard Ng, Devin M. Saunders, Amiko K. Lagrimas, Duc Duong, Nicholas T. Seyfried, David A. Bennett, Yanling Wang

## Abstract

The choroid plexus (CP) regulates barrier integrity, cerebrospinal-fluid (CSF) dynamics, and immune surveillance, yet its role in Alzheimer’s disease (AD) remains poorly defined. We performed snRNA-seq on CP samples from 69 ROSMAP participants spanning normal cognition, mild cognitive impairment, and AD dementia, and integrated these data with spatial transcriptomics, snATAC-seq, and proteomics from CP tissue and CSF. We identified 17 CP cell states and uncovered widespread disease-associated transitions that converged into three major phenotypic axes. Along the inflammatory axis, epithelial cells and border-associated macrophages (BAMs) showed progressive immune activation, with BAMs shifting from inflammatory to stress-dominant states. In the barrier axis, epithelial cells, fibroblasts, and endothelial cells exhibited reduced junction-related gene expression and broad alterations in transport pathways. Epithelial cells also showed late-stage cilia loss and CSF-regulatory pathway impairment, indicating a breakdown in epithelial polarity and CSF sensing, consistent with abnormal CSF proteomic signatures. Along the remodeling axis, fibroblasts showed ECM alterations, while epithelial and stromal cells demonstrated aberrant cell–matrix adhesion pathways. Spatial neighborhood analysis revealed a multicellular signaling hub, with epithelial-rich niches showing the strongest dysregulation in AD. Together, these findings define a unified model of CP dysfunction in AD and position the CP as an active, multicellular contributor to AD pathophysiology.

## Introduction

The choroid plexus (CP) has traditionally been viewed as a supporting structure that forms the interface between the brain and periphery, producing cerebrospinal fluid (CSF). However, emerging evidence suggests that the CP is a dynamic regulator of brain homeostasis, modulating barrier permeability, molecular transport, metabolism, and immune surveillance^1–4^. Disruption of CP function has been implicated in impaired CSF flow, barrier breakdown, and neuroinflammation—processes that are highly relevant to Alzheimer’s disease (AD)^5–7^.

Despite these links, the CP has received considerably less attention in AD research than the brain parenchyma. Studies using animal models or human tissue suggest that CP barrier integrity and function are altered in AD, potentially affecting immune cell trafficking and the clearance of pathological proteins^8–15^. However, a comprehensive understanding of how human CP cell states shift along the clinical continuum from no cognitive impairment (NCI) to mild cognitive impairment (MCI) and Alzheimer’s dementia (ADd) remains lacking.

To address this gap, we applied high-resolution single-cell multi-omics and spatial transcriptomics to construct a detailed atlas of CP alterations across NCI, MCI, and ADd individuals from the Religious Orders Study or Rush Memory and Aging Project (ROSMAP)^16^. Our analysis reveals coordinated cell state differences in epithelial, immune, endothelial, and fibroblast populations that collectively reshape the CP’s immune, barrier, and stromal landscape during disease progression, offering new mechanistic insights into how CP dysfunction contributes to AD pathophysiology.

## Results

### Major cell classes of the human choroid plexus

To transcriptionally characterize the CP cell types and states, we performed snRNA-seq on 69 individuals (NCI=21, MCI=21, and ADd=27; Fig. 1a-b; Extended Data table 1). After quality control and doublet removal, we recovered 758,302 total nuclei. Based on canonical markers, we annotated cell clusters and detected five major CP cell types, including epithelial (*HTR2C, CLIC6, CFAP54, DNAH11*), fibroblast (*DCN*, *LEPR*, *FN1*), endothelial (*PECAM1*, *VWF*, *INSR*), mural (*ACTA2*, *TAGLN*, *MYH11*), and immune (*CD163*, *STAB1*, *MSR1*) populations (Fig. 1c–d, Extended Data table 2). Gene detection per nucleus was consistent with previous postmortem human snRNA-seq studies^18–20^ (Extended Data Fig. 1a-c), and no significant cell-type proportion differences were observed between NCI, MCI and ADd groups (Fig. 1e Extended Data table 3).

**Fig. 1:**
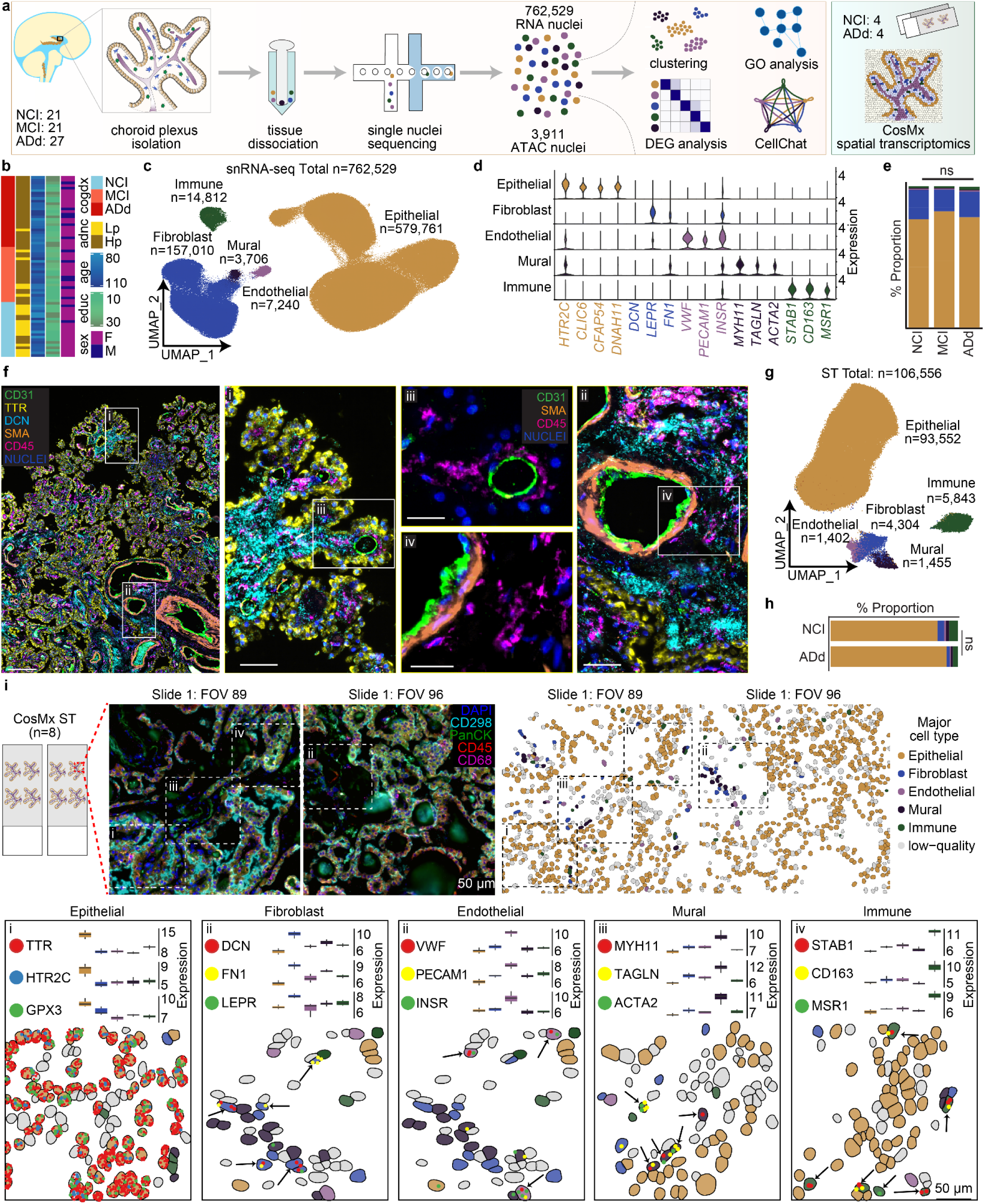
Major cell classes of the human choroid plexus. **a,** Study design overview. snRNA-seq was performed on 69 participants (762,529 nuclei) and snATAC-seq was performed on nuclei from 6 participants (3,238 nuclei), and spatial transcriptomics (ST) on 8 participants (106,556 segmented cells). **b,** Participant demographics, balanced for age, sex, and education (educ). Clinical (cogdx) and neuropathological (ad_adnc) diagnoses are indicated. See Extended Data Table 1. **c,** snRNA-seq UMAP colored by major cell types. **d,** snRNA-seq canonical marker expression violin plot per cell cluster (pseudobulk FDR < 0.01, log₂FC > 0.25). See Extended Data Table 2. **e,** snRNA-seq major cell type proportions stratified by clinical diagnosis (all not significant, FDR > 0.01, ns). **f,** Multiplexed immunohistochemistry staining for CD31, TTR, DCN, SMA, CD45 and DAPI. Scale bars = 200 µm (overview), 50 µm (i, ii) and 20 µm (iii, iv). **g,** Unsupervised spatial transcriptomics (ST) UMAP projection of major cell types. **h,** Proportion of epithelial subclusters in ST data. **i,** Adjacent spatial transcriptomics fields of view (FOVs #1.89 and #1.96) Upper left: immunostaining. Upper right: segmented cells colored by major cell type. Below: transcript localization with quantified pseudobulk expression (log_2_counts; upper) and example FOV (lower). Arrows indicate marker localization. Scale bars, 50 µm.

We next validated the major CP cell types by multiplex immunohistochemistry. Consistent with previous human histological studies^21^, we detected TTR⁺ epithelial cells forming the luminal boundary, DCN⁺ fibroblasts enriched in stromal regions, CD31/PECAM1⁺ endothelial cells comprising small and large vessels, SMA⁺ mural cells constituting the large vessel wall, and CD45⁺ immune cells dispersed across stromal and epithelial compartments (Fig. 1f). Together, these results confirm the identity of major CP cell types and highlight the widespread presence of immune cells across stromal and vascular compartments.

To validate marker gene expression and spatially contextualize these findings, we applied spatial gene profiling using the CosMx Spatial Molecular Imager on CP sections from eight individuals (NCI=4; AD=4; Extended Data table 1, Fig 1i, upper left) with the 6K human discovery panel. After excluding low-quality or mis-segmented cells, we obtained 106,556 high-quality cells for analysis. Unbiased clustering and annotation revealed the same major CP cell types identified in snRNA-seq (Fig. 1g, Extended Data Fig 1d-i, Extended Data table 4), with no significant differences in cell-type proportions between NCI and AD (Fig. 1h, Extended Data table 5). We confirmed the expected localization of annotated cell types, including epithelial, fibroblast, endothelial, mural, and immune cells (Fig. 1i, top). At the gene level, canonical markers were enriched in their expected cell types: *TTR*, *HTR2C*, and *GPX3* in epithelial cells; *DCN*, *FN1*, and *LEPR* in fibroblasts; *VWF*, *PECAM1*, and *INSR* in endothelial cells; *MYH11*, *TAGLN*, and *ACTA2* in mural cells; and *STAB1*, *CD163*, and *MSR1* in immune cells (Fig. 1i, bottom). These data validate cell-type assignments and demonstrate the spatial organization of CP cellular compartments.

We conducted multi-omic profiling (snATAC-seq with matched snRNA-seq) in a subset of cases (n=6) to assess chromatin accessibility alongside gene expression. After standard QC, UMAP clustering identified major epithelial, fibroblast and immune clusters, with ATAC accessibility patterns concordant with RNA-based cell type assignments (Extended Data Fig. k-q, Extended Data table 6).

### Epithelial cells exhibit progressive inflammatory activation and late ciliary dysfunction in the choroid plexus across disease stages

We subclustered CP epithelial cells into four transcriptionally distinct epithelial subtypes: Epi_1a, Epi_1b, Epi_2a, and Epi_2b (Fig. 2a-e, Extended Data Fig. 2a-d, and tables 7-9). Epi_2a/2b were enriched for transport and secretory genes (*TTR*, *AQP1*, *SLC25A6*, *SLC25A23*, *SLC4A2*), and Epi_1a/1b/2a were enriched for ciliogenesis signatures (*DNAH6*, *CFAP100*, *CFAP69*). Epi_2a/2b showed higher tight-junction proteins (*CLDN1/5*, *OCLN*) and greater mitochondrial gene expression (*MT-CO1*, *MT-CO2*, *MT-CO3*) (Fig. 2b, Extended Data table 7). Functional annotation further aligned with marker gene profiles: Epi_1a/1b and Epi_1b/2a were associated with microtubule movement and cilium assembly, and Epi_2b was enriched for transport and metabolic processes (Fig. 2c).

**Fig. 2:**
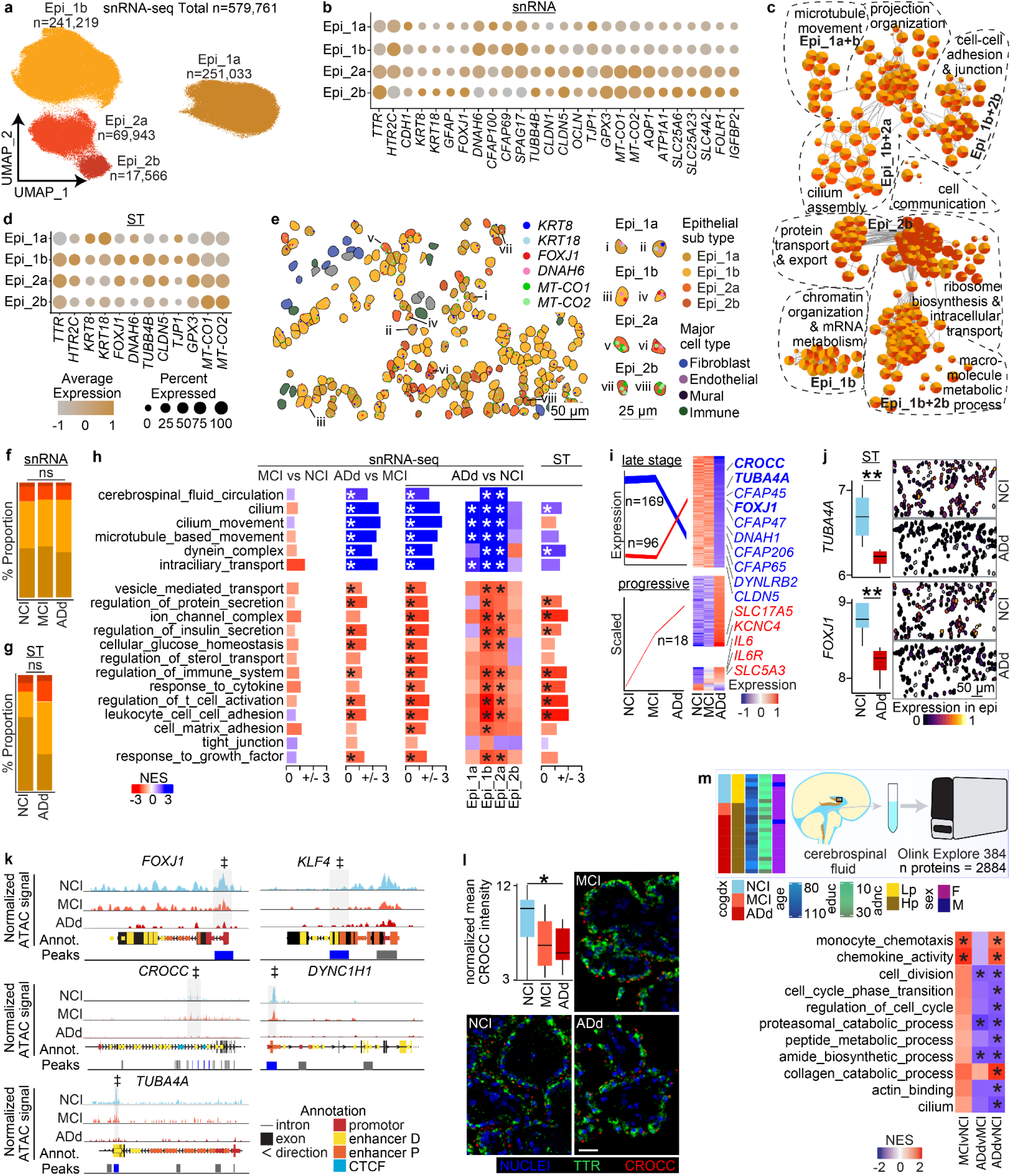
Progressive epithelial inflammation and late ciliary loss in Alzheimer’s disease. **a,** snRNA-seq UMAP projection colored by epithelial subtypes. **b,** Epithelial subtype-defining marker dot plot (FDR < 0.01, log₂FC > 0.25). See Extended Data Table 7. **c,** Epithelial subtype GO enrichment network plot (ClueGO on subtype-defining markers) grouped by functional category. **d,** Unsupervised spatial transcriptomics (ST) confirmation of epithelial subtype-defining marker dot plot. See Extended Data Table 8. **e,** Spatial localization of epithelial subtypes overlaid with subtype-defining transcript localization; FOV: 1.186. Scale bar: 50µm. Right: Selected examples. Scale bar: 25 µm. **f-g,** snRNA-seq (f) and spatial transcriptomics (g) epithelial subtype proportions across disease states. Linear mixed model on proportions (* FDR < 0.01). **h,** Select GO pathways (via fgsea on ranked t statistics) of DEGs throughout clinical disease in snRNA-seq (SN) and matched spatial transcriptomics (ST) (*FDR < 0.15). See Extended Data Table 10-12. **i,** Disease temporal dynamics of DEGs. “Late-stage specific” genes: ADd ≠ NCI & MCI; “progressive” genes: NCI ≠ MCI & ADd and MCI ≠ ADd. Left: line plots indicate the mean of scaled gene expression; line width corresponds to the number of genes following each trajectory. Right: heatmap per stage select genes representing key GO pathways are labelled (nominal p < 0.05, log₂FC > ±0.25, trajectory correlation p < 0.05). **j,** Spatial transcriptomics expression of select cilia genes (left: boxplots of pseudobulk expression (log_2_counts), median, interquartile range, min/max, ** FDR < 0.01; right: normalized expression in epithelial cells in example FOVs for NCI or ADd). Scale bars, 50 µm. **k**, Differentially accessible chromatin peaks of select cilia genes throughout clinical disease progression (‡ nominal p<0.05 ADd vs NCI). Genome-specific annotations and accessible peaks are shown below. See Extended Data Table 13. **l**, Mean CROCC intensity per TTR+ epithelial cell normalized to Hoechst intensity. Upper left: box plots throughout clinical disease progression (median, interquartile range, min/max; n = 9 NCI, 12 MCI, 9 ADd). One-way ANOVA F (2,27) =4.135, p<0.05; Tukey’s HSD * p<0.05. Right/lower: representative immunohistochemistry of DAPI, TTR and CROCC per group. Scale bar: 50 µm. **m,** Selected GO pathways of CSF protein data throughout clinical disease progression (via fgsea on ranked t statistics; * FDR < 0.15). See Extended Data Table 14.

Unsupervised clustering across spatial transcriptomics (ST) and ATAC-seq consistently identified four major epithelial subclusters (Fig. 2d, Extended data Table 8; Extended Data Fig. 2a-d and Table 8-9). Although individual marker genes showed variable assignments across data types, likely reflecting modality-specific resolution, we identified both ciliated and less-ciliated cell states within epithelial cells (Fig. 2d–e, Extended Data Table 8-9; Extended Data Fig. 2a-d).

Importantly, the relative proportions of the four epithelial subclusters were stable across NCI, MCI, and ADd in our snRNA-seq and ST data (Fig. 2f-g, Extended Data Table 3 and 5), indicating that epithelial heterogeneity itself is not disease-associated. We next examined transcriptome differences across AD disease progression at both the aggregated epithelial cell level and within individual epithelial subtypes. Along the disease trajectory from NCI to MCI to ADd, we detected progressive upregulation of immune and inflammatory genes (e.g. *IL6R* and *SERPINE3*). However, we observed a marked downregulation of cilia-related genes (e.g. *CROCC*, *FOXJ1*, *TUBA1A*, and *CFAP43*) that correlated with cognitive decline (Extended Data Fig. 2e and Table 10-11). Gene set enrichment analysis revealed that immune responses, secretion, and transporter activities were progressively higher from NCI to ADd, whereas cilia-related pathways were selectively lower in ADd (Fig. 2h). At the epithelial subtype level, Epi_1b/2a carried the strongest signatures of both immune activation and cilia loss (Fig. 2h). ST corroborated these findings at the aggregated epithelial cell level (Fig 2h right and Extended Data Table 12).

To resolve disease temporal patterns, we used template matching^22^ to classify disease-associated transcriptional programs into different patterns and identified two major categories: progressive differences included immune and transporter genes (e.g., *SLC17A5*, *IL6R*; Fig. 2i), while late-stage changes were dominated by cilia-related genes (*CROCC*, *FOXJ1*, *TUBA4A*; Fig. 2i and Extended Data Table 10). ST corroborated these findings, revealing reduced cilia gene expression (*FOXJ1*, *TUBA4A*) in ADd epithelium (Fig. 2j). Chromatin accessibility profiling by snATAC-seq further validated these observations, showing less accessibility at key cilia-related transcription factor loci (e.g. *FOXJ1, KLF4)*) and correspondingly structural gene loci (*CROCC, DYNC1H1* and *TUBA4A*), consistent with their transcriptional downregulation (Fig. 2k, Extended Data Table 13). Immunohistochemistry confirmed a reduction of ciliary rootlets, with CROCC staining markedly lower in AD relative to controls (Fig. 2l).

Since the CP epithelium is a key tissue interface with CSF, we next examined protein expression in CSF using the Olink Explore platform (n=27), which enabled measurement of 2,884 proteins (Fig. 2m, upper). This analysis revealed stage-dependent differences that paralleled the transcriptomic findings. At the early stage of AD progression, we observed more inflammatory signatures, including enrichment of monocyte chemotaxis and chemokine activity pathways (Fig. 2m). By the late stage, we detected lower cilium-related terms and multiple changes to metabolic processes, including proteasomal and collagen catabolism (Fig. 2m; Extended Data Table 14). Together, these data highlight coordinated molecular changes between CP epithelial cells and the CSF proteome, reinforcing the role of the CP as a dynamic interface linking inflammation and impaired CSF secretion mechanisms in AD.

### Border-associated macrophages shifted from inflammatory to stress-associated states in the choroid plexus across disease stages

We next examined immune cells in the CP. Multiplex fluorescent imaging confirmed the presence of CD163⁺ macrophages and CD3⁺ T cells within the ECAD+ CP tissue (Fig. 3a). Subclustering of snRNA-seq data identified three states of border-associated macrophages (BAM_1-3) and a small T-cell population marked by expression of *THEMIS* and *CD3E* (Fig. 3b). BAM_1 expressed *CD163*, *MRC1*, *LYVE1*, and *IL4R*, consistent with an immune-activated state. BAM_2 was enriched for heat-shock protein genes (*HSPE1*, *-*, *HSPA8, DNAJA1*), indicative of a stress-adapted phenotype, while BAM_3 expressed *MKI67* and *CENPF*, reflecting a proliferative subset. Functional enrichment supported these distinctions: BAM_1 was associated with immune activation and cytokine release, BAM_2 with protein folding and stress responses, and BAM_3 with cell cycle and proliferation (Fig. 3c-d, Extended Data Table 15). snATAC-seq revealed selective chromatin accessibility at *DNAJA1* and *HSPA8* loci within the BAM2 subcluster, consistent with their elevated transcription (Fig. 3e, Extended Data Fig. 3a-b and Table 16). Notably, BAM1- and BAM2-specific transcriptional signatures corresponded with distinct ATAC activity, indicating coordinated regulation of chromatin accessibility and gene expression across BAM states (Fig. 3f). Because of limited immune cell numbers, we applied a supervised approach^23^ to annotate ST subclusters, which showed strong correspondence with snRNA-seq–defined BAM populations (Fig. 3g-h, Extended data Fig 3c). When mapped back to the CP, BAM1, BAM2, BAM3, and T cells were broadly distributed, confirming the presence of these immune populations throughout the tissue (Fig. 3h).

**Fig. 3:**
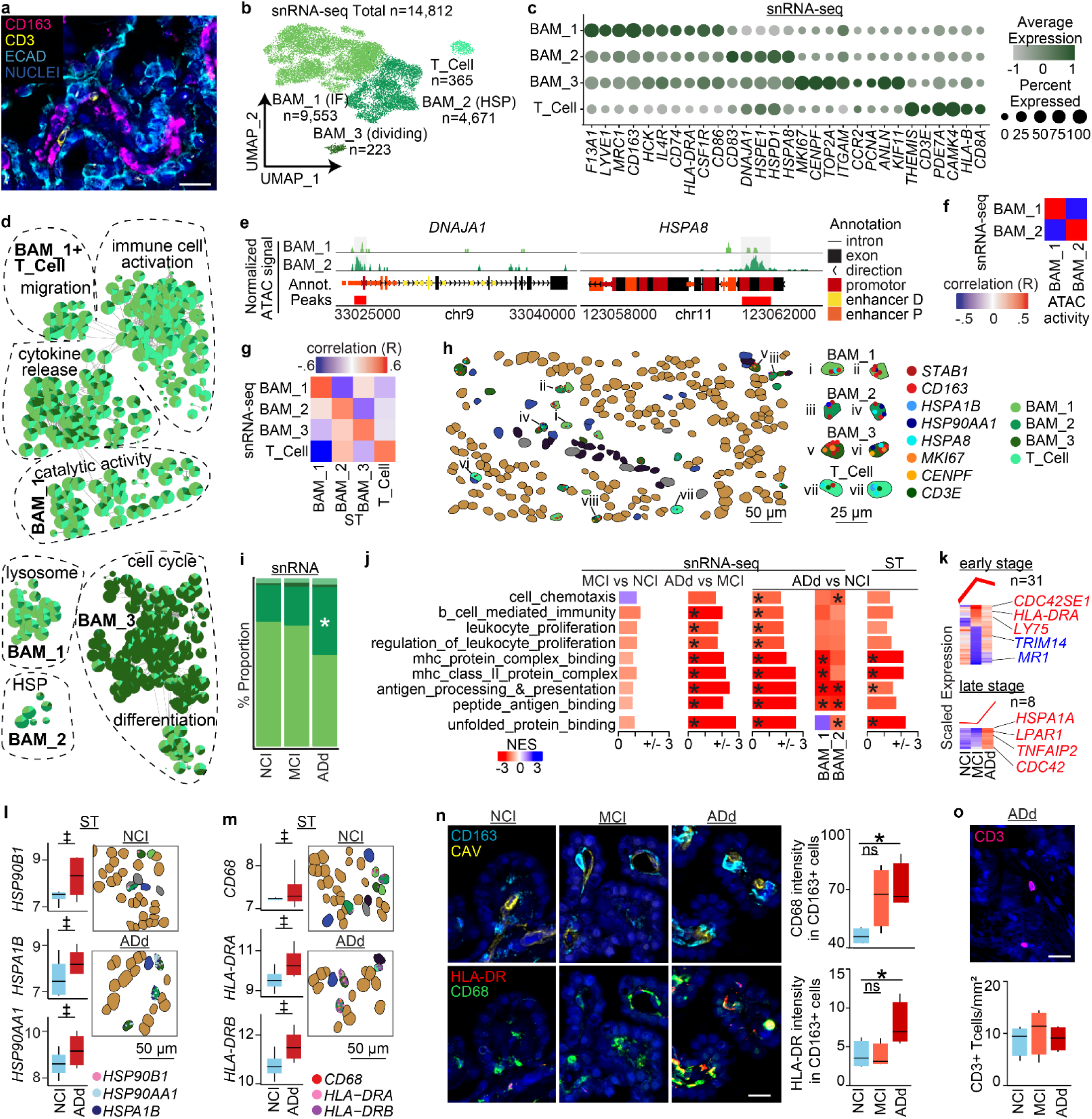
Border-associated macrophages shifted from inflammatory to stress-associated states in Alzheimer’s disease. **a,** Phenocycler fusion multiplexed immunostaining of DAPI, CD163, CD3 and E-cadherin (ECAD). Scale bar: 50µm. **b,** snRNA-seq UMAP of immune cell subclusters. **c,** Immune subtype defining marker dot plot (FDR < 0.01, log₂ fold change > 0.25). See Table Extended Data Table 15. **d,** Immune subtype GO enrichment network plot (ClueGo on subtype-defining markers) grouped by functional category and color-coded by immune subtype. **e,** Differentially accessible chromatin peaks of select HSP genes per subtype. Genome-specific annotations and accessible peaks are shown below. **f,** Pearson correlation of chromatin accessibility (ATAC activity) and snRNA-seq expression among shared immune substates. **g,** Pearson correlation of supervised spatial transcriptomics (ST) and snRNA-seq expression among immune substates. **h,** Spatial localization of immune subtypes overlaid with subtype-defining marker expression; FOV: 1.186. Scale bar: 50µm. Right: Selected examples. Scale bar: 25 µm. **i,** Subtype proportions stratified by clinical diagnosis (* BAM_2 FDR < 0.01). **j,** Select GO pathways (via fgsea on ranked t statistics) of DEGs throughout clinical disease in snRNA-seq and matched spatial transcriptomics (*FDR < 0.15). See Extended Data Table 17-19. **k,** Disease temporal dynamics of DEGs. “early stage” genes: NCI vs. MCI; “Late-stage specific” genes: ADd ≠ NCI & MCI. Upper: line plots indicate the mean of scaled gene expression; line width corresponds to the number of genes following each trajectory. Lower: heatmap per stage; select genes representing key GO pathways are labelled (p < 0.05, log₂FC > ±0.25, trajectory correlation p < 0.05). **l-m,** Spatial transcriptomics gene expression of select HSP (**l**) and inflammation (**m**) Left: boxplots of pseudobulk expression (log_2_counts) (median, interquartile range, min/max, ‡ nominal p < 0.05). Right: Spatial localization of expression in NCI FOV #1.168 and AD FOV #2.24. Scale bar: 50 µm. **n,** Left: Representative Phenocycler fusion immunofluorescence of HLA-DR and CD68, or CD163 and caveolin (CAV) with DAPI for NCI and ADd. Scale bar: 50µm. Right: box plots of mean HLA-DR or CD68 intensity within CD163+ cells in NCI or ADd (median, interquartile range, and full range, *p ≤ 0.05). Welch’s test HLA-DR: p=0.0582; CD68: p=0.01. **o,** Representative Phenocycler fusion immunofluorescence of DAPI, CD3 and CD163. Scale bar, 50µm. Right: Box plot of the numbers of CD4+ and CD8+ CD3+ T cells in NCI or ADd tissue (median, interquartile range, and full range). Non-significant (ns).

Notably, we observed higher BAM_2 proportions in ADd (Fig. 3i, Extended Data Table 3), indicating a shifted state toward a stress-dominant phenotype. At the pathway level, we observed progressively stronger immune responses across disease stages, including enrichment of pathways such as antigen presentation, leukocyte proliferation, and other signatures of immune activation. In ADd, we also detected enrichment of unfolded protein binding, indicative of stress responses, consistent with the expansion of the BAM_2 population characterized by high expression of heat shock proteins (Fig. 3j). At the gene level, we observed stage-dependent regulation of classical immune-related genes, including *HLA-DRA*, as well as additional immune-associated genes linked to CP function, such as *CDC42SE1*, *LY75*, *TRIM14*, and *MR1* in early stages, and *LPAR1*, *TNFAIP2*, *CDC42*, and *HSPA1*A in late stages (Fig. 3k, Extended Data Table 17). Spatial transcriptomics data confirmed increased *HSP* expression, along with elevated *CD68* and *HLA-DRA/DRB* expression in AD (Fig. 3j, right and l-m, Extended Data Table 18). Immunostaining further validated these findings, showing higher HLA-DR and CD68 intensity in CD163⁺ macrophages in AD compared with NCI (Fig. 3n). However, we did not observe significant differences in the number of total T cells, CD4⁺ T cells, or CD8⁺ cells in AD (Fig. 3o, Extended Data Fig. 3d). Together, these results demonstrate that immune cells in the CP are heterogeneous and undergo progressive immune activation and stress responses across disease stages.

### Fibroblast remodeling in the choroid plexus in AD dementia

To characterize fibroblast heterogeneity in the CP, we subclustered fibroblasts into three transcriptional subtypes (Fib_1, Fib_2, Fib_3; Fig. 4a). Fib_1 expressed markers such as *LEPR*, *PLXNA4*, and *STXBP6*, consistent with regulatory and signaling functions. Fib_2 was enriched for *CLDN11*, *TJP1, ITGBL1, GJB6, TRPM3*, and *ROBO1*, indicative of tight-junction-expressing barrier-like fibroblasts. Fib_3 expressed *DCN*, *FN1*, *FBLN1*, *CDH11*, and multiple collagens (e.g., *COL4A2*, *COL3A1*, *COL6A3*), indicative of extracellular-matrix (ECM)-producing fibroblasts (Fig. 4b, Extended Data Table 19). Recent work identified a population of “base barrier” fibroblasts in the mouse CP using scRNA-seq^2^, describing them as a barrier-forming population at the stroma–CSF interface. While similar gene signatures were noted in a published human dataset^19^, those base barrier fibroblasts had not been independently resolved in human tissue due to limited sampling. Our study extends these findings using unbiased clustering of a larger cohort, resolving Fibro_2 as the human counterpart of this base barrier fibroblast population. Immunohistochemistry staining confirmed that CLDN11⁺ Fibro_2 localized to the basal side of the CP epithelium, particularly around larger vessels (CD31, SMA) adjacent to the brain parenchyma, consistent with a barrier-supporting niche (Fig. 4c).

**Fig. 4:**
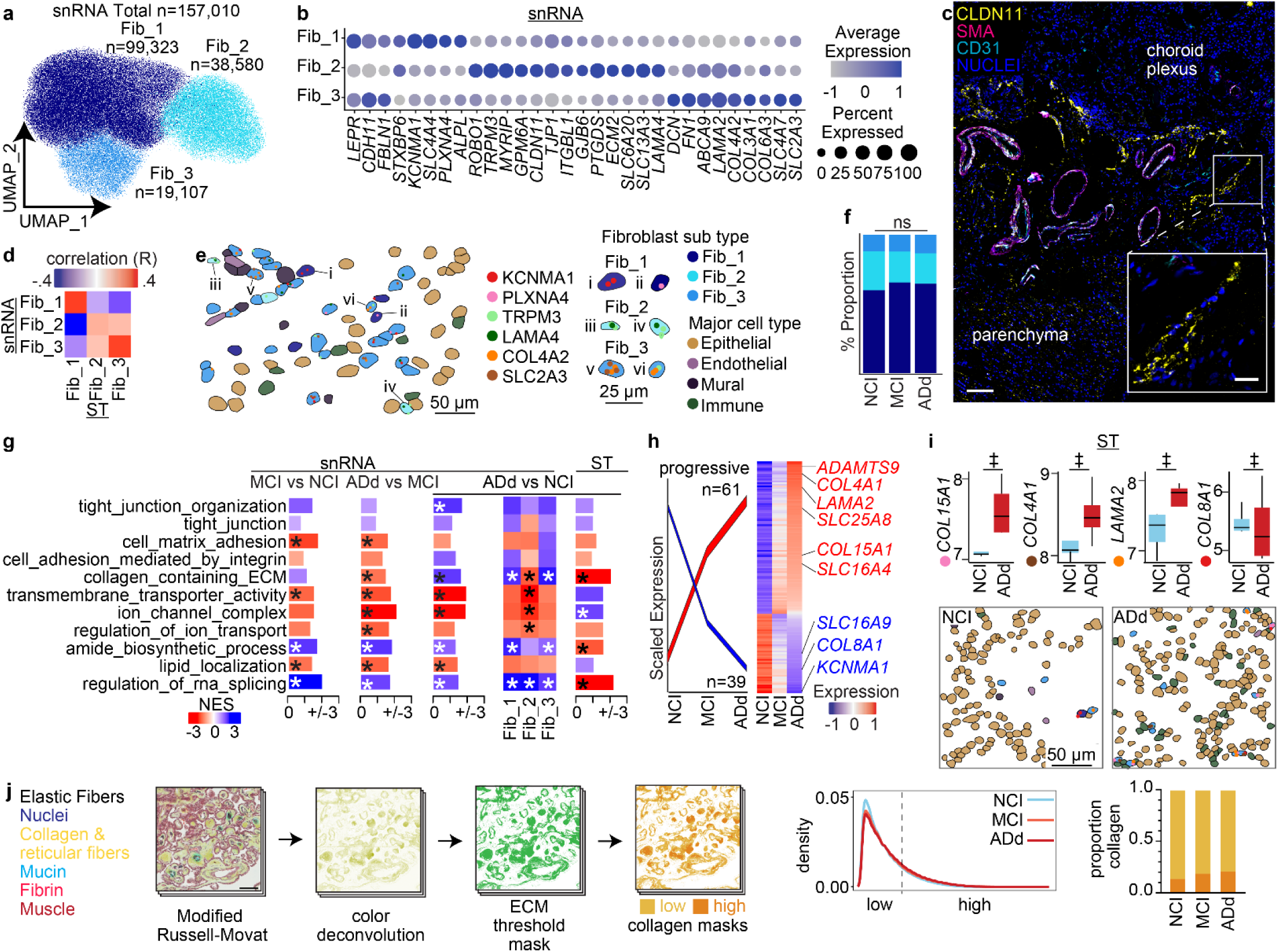
Fibroblast subtypes and remodeling in Alzheimer’s Disease. **a**, snRNA-seq UMAP projection colored by fibroblast subtypes. **b**, Fibroblast subtype-defining marker dot plot (FDR < 0.01, log₂FC > 0.25). See Extended Data Table 19. **c**, Multiplexed Opal immunohistochemistry for DAPI, CLDN11, CD31 and SMA, showing localization of CLDN11-positive cells at the boundary between CP and parenchyma. Scale bars = 100µm (overview), 20µm (inset). **d,** Pearson correlation of supervised spatial transcriptomics (ST) and snRNA-seq expression among fibroblast subtypes. **e**, Spatial localization of fibroblast subtypes overlaid with subtype-defining marker expression; FOV: 2.160. Scale bar: 50µm. Higher magnification on right. Right: Selected examples. Scale bar: 25 µm. **f,** Subtype proportions stratified by clinical diagnosis Linear mixed model on proportions (all not significant, FDR > 0.01, ns). **g**, Select GO pathways (via fgsea on ranked t statistics) of DEGs throughout clinical disease in snRNA-seq and matched spatial transcriptomics (* FDR < 0.15). See Extended Data Table 20-22. **h**, Disease temporal dynamics of DEGs. “progressive” genes: NCI ≠ MCI & ADd and MCI ≠ ADd. Left: line plots indicate the mean of scaled gene expression; line width corresponds to the number of genes following each trajectory. Right: heatmap per stage select genes representing key GO pathways are labelled (nominal p < 0.05, log₂FC > ±0.25, trajectory correlation p < 0.05). **i,** Spatial transcriptomic expression of select ECM genes. Upper: boxplots of pseudobulk expression (log_2_counts) (median, interquartile range, min/max, **‡** nominal p < 0.05); bottom: transcript localization in fibroblasts in NCI FOV #1.86 and ADd FOV #2.119. Scale bar: 50 µm. **j**, Upper and lower left: Representative Modified Russel-Movat’s stain, deconvoluted yellow (collagen), and ECM threshold mask (< 230 pixel intensity), and high and low collagen masks (230-200, 200-0) (Scale bar: 100µm). Lower middle: averaged distribution of collagen intensity colored by disease stage. Lower right: Stacked barchart of the proportion of high and low collagen by disease stage. n = 8 NCI, 10 MCI, 9 ADd, 5-10 FOVs per subject

Despite these molecular distinctions, gene ontology enrichment showed broad overlap across fibroblast subtypes, with shared programs in adhesion, ECM organization, and ion transport (Extended Data Fig. 4a).We then applied the same supervised approach^23^ to annotate ST fibroblast subclusters, which showed modest correspondence with snRNA-seq–defined populations (Extended Data Fig. 4b; Fig. 4d) and mapped to original locations within the CP (Fig. 4e).

We next examined disease-associated transcriptional differences at the fibroblast subtype level. Fibroblast proportions were similar across NCI, MCI, and ADd (Fig. 4f), indicating that disease progression does not alter relative abundance but instead drives transcriptional reprogramming within each subtype. Gene set enrichment analyses revealed broad transcriptional reprogramming across major pathways, with changes most pronounced at the late stage (Fig. 4g and Extended Data Table 20-21). Tight junction-related pathways showed overall negative enrichment, indicating weakened barrier integrity. By contrast, pathways related to cell–matrix adhesion, transporter, and ion channel pathways displayed positive enrichment at the late stage, indicating an adaptive transcriptional response. Among fibroblast subtypes, Fibro_2 showed the most consistent upregulation of ECM- and transporter-related pathways. At the tissue level, ST data revealed a general downregulation of transporter and ion channel activity and upregulation of ECM remodeling. We observed that pathway directionality is not always concordant between snRNA-seq and ST, likely reflecting differences in cell-type purity, spatial mixing within ST spots, and modality-specific sensitivity to fibroblast state composition. Together, these findings indicate that fibroblast remodeling in ADd involves broad transcriptional changes, with consistent tight junction weakening and ECM remodeling.

To further resolve temporal differences, we examined gene expression patterns in fibroblasts. Several collagen-related genes, including *ADAMTS9*, *COL15A1*, and *LAMA2*, were progressively upregulated, consistent with greater ECM deposition, whereas others, such as *COL8A1*, were downregulated, indicating selective collagen loss (Fig. 4h Extended Data Fig. 4c). Similarly, transporter genes, such as the SLC family (*SLC25A8*, *SLC16A4*, *SLC16A9*), also exhibited bidirectional regulation, highlighting heterogeneous shifts in ion and metabolite handling. To examine stage-dependent transcriptional dynamics, we compared differentially expressed genes (DEGs) across fibroblast subtypes at each disease stage. The overlap between DEGs across stages was mild to moderate, suggesting progressive but partly distinct transcriptional alterations during disease progression. Similarly, when comparing ADd versus NCI across different fibroblast subtypes, only limited overlap was observed, indicating that each fibroblast subtype exhibits partially unique transcriptional responses to AD pathology (Extended Data Fig. 4d-e).

ST analysis confirmed these fibroblast-specific transcriptional changes in AD, with increased *ADAMTS9*, *COL15A1*, and *LAMA2* expression and reduced *COL8A1* relative to controls (Fig. 4i). To validate these findings at the histological level, we performed Movat’s pentachrome staining, followed by color deconvolution and ECM threshold masking to quantify collagen deposition (Fig. 4j). The density distribution analysis revealed a subtle shift from low- to high-collagen regions in AD and MCI compared with NCI although this trend did not reach statistical significance (Fig. 4j).

### Endothelial and mural cell dysregulation in the choroid plexus in AD dementia

We next investigated vascular compartments of the CP, focusing on endothelial and mural cells. We identified three endothelial subtypes (Endo_1, Endo_2, Endo_3) and three mural subtypes (Mural_1, Mural_2, Mural_3. Endo_1 was characterized by strong *PECAM1*, *CLDN5*, and *CDH13* expression with lower *VWF*, consistent with an arterial/arteriolar endothelial identity. Endo_2 also expressed *CLDN5* but showed higher *VWF* together with *CEMIP2*, *IL1R1*, and *IL4R*, consistent with a venous endothelial phenotype. Endo_3 was distinguished by robust *PLVAP* and *FLT1* expression, consistent with fenestrated capillary endothelium in the CP (Fig 5a-b, Extended Data Table 22). Mural_1 expressed high levels of canonical contractile smooth muscle markers, including *ACTA2*, *MYH11*, *TPM1*, and *MYOCD*, consistent with arterial smooth muscle cells. Mural_2 showed lower expression of contractile genes but enrichment of ECM-remodeling transcripts such as *COL18A1*, *ADAMTS9*, and *CRISPLD2*, indicative of a venous smooth muscle or transitional phenotype. Mural_3 exhibited reduced expression of smooth muscle genes and upregulation of signaling and matrix regulators such as *CEMIP*, *PLXDC2*, and *ENPP2*, consistent with a pericyte identity associated with capillaries (Fig 5c-d, Extended Data Table 23). We then applied the same supervised approach^23^ to identify ST vascular subtypes, which showed strong correspondence with snRNA-seq–defined populations (Fig. 5e) and mapped to vascular niches within the CP (Fig. 5g).

**Fig. 5:**
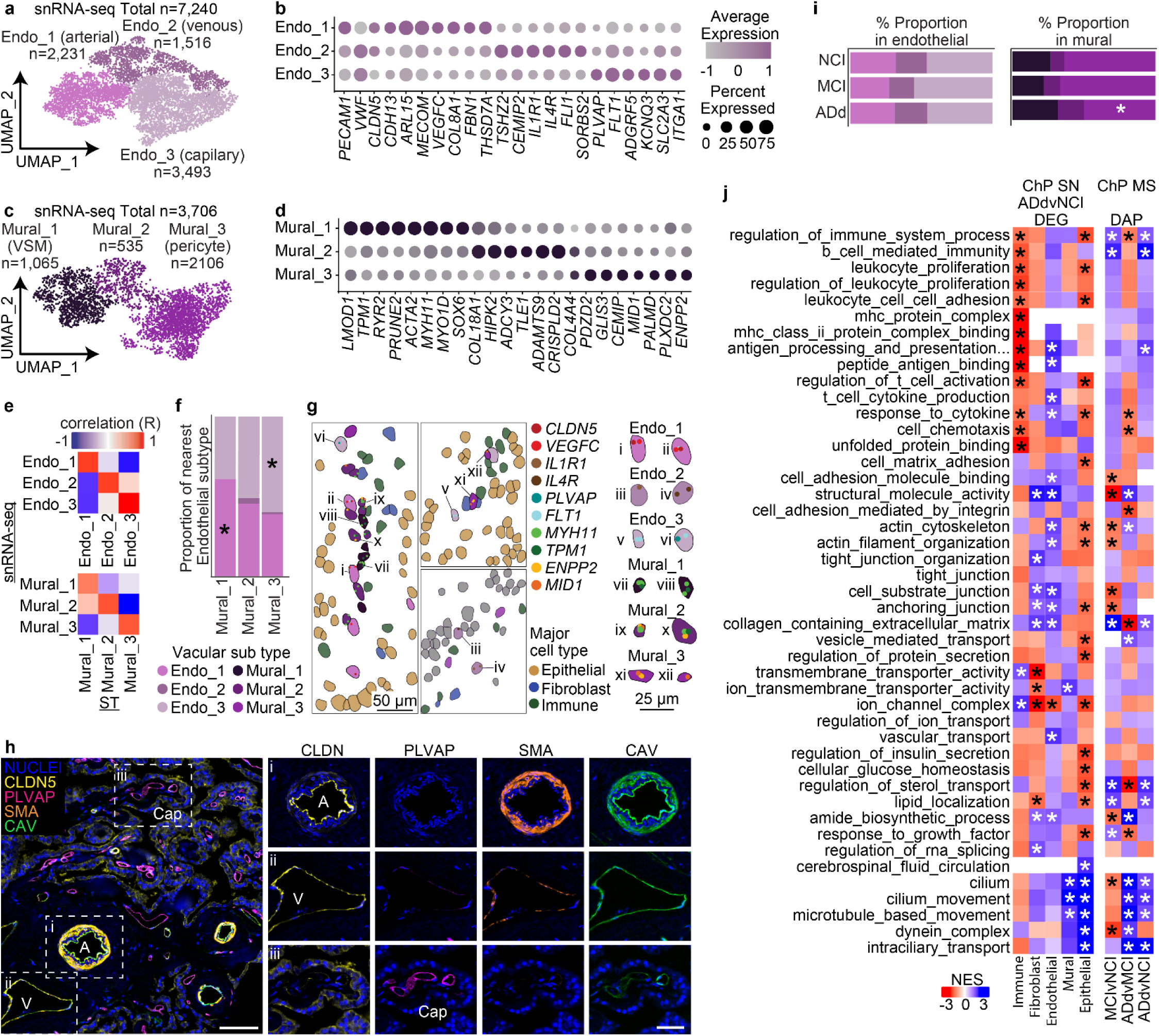
Endothelial and mural cell dysregulation in Alzheimer’s Disease. **a, & c,** snRNA-seq UMAP colored by endothelial (a) and mural (c) subtypes. **b, & d,** Endothelial (b) and mural (d) subtype-defining marker dot plot (FDR < 0.01, log₂FC > 0.25). See Extended Data Tables 22-23. **e,** Pearson correlation of supervised spatial transcriptomics (ST) to snRNA-seq expression pooled among endothelial (upper) and mural (lower) subtypes. **f**, Proportion of nearest endothelial subtype across Mural subtypes. Linear mixed model on proportions (* p < 0.05). **g,** Spatial localization of endothelial and mural subtypes overlaid with subtype-defining marker expression; FOV: 1.186; Scale bar: 50µm. Right: Selected examples. Scale bar: 25 µm. **h**, Phenocycler fusion immunofluorescence of endothelial (CLDN5, CAV, PLVAP) and mural (SMA) with DAPI. Capillary (Cap), arterial (A), and vascular venous(V) structures indicated and split by stain on right. **i,** Endothelial (left) and mural (right) subtype proportions stratified by clinical diagnosis. Scale bars = 100µm (overview), 50µm (insets); Mural_3 * FDR < 0.01, others not significant. **j**, Select GO pathways (via fgsea on ranked t statistics) of ADd vs NCI DEGs across all major cell types matched to mass spectrophotometry proteomics throughout clinical disease (*FDR < 0.15). See Extended Data Tables 24-28.

We next asked how mural subtypes are positioned relative to endothelial populations (Fig. 5f). Mural_1, which we annotated as arterial smooth muscle cells, was preferentially associated with Endo_1 (arterial endothelium). By contrast, Mural_3, defined as pericytes, showed enrichment near Endo_3, corresponding to fenestrated capillary endothelium. These relationships indicate coordinated arterial and capillary compartmentalization between endothelial and mural cells in the choroid plexus.

To validate the endothelial subtype classification, we performed immunostaining with CLDN5 and PLVAP together with SMA and CAV to visualize vascular structure (Fig. 5h). CLDN5 was strongly expressed in arteries and veins, whereas PLVAP was selectively enriched in capillaries. Arteries and veins were further distinguished by mural SMA staining, with arteries showing thick, robust expression and veins exhibiting thinner signals. These findings confirmed that CLDN5 and PLVAP are reliable markers for distinguishing arterial/venous from capillary endothelial cells. The endothelial cell representation did not differ across NCI, MCI, and ADd groups, whereas mural populations had a lower Mural_3 subtype in ADd (Fig. 5i). Given the pericyte-like transcriptional profile of Mural_3, this decrease may reflect pericyte loss or dedifferentiation, indicating vascular instability and barrier dysfunction in ADd. Correspondingly, differential expression analyses in snRNA-seq and ST revealed progressive reductions in junctional and actin-binding pathways, along with downregulation of ion transport signatures, consistent with progressive barrier weakening in ADd (Extended Data Tables 24-27).

### Transcriptomic–proteomic concordance highlights multicellular dysfunction in the choroid plexus in AD dementia

To determine whether cell type–specific transcriptomic changes are mirrored at the protein level, we compared snRNA-seq–derived differential gene signatures (ADd vs. NCI) across major cell types with bulk CP proteomic profiles stratified by disease stage (Fig. 5j, Extended Data Table 28). Transcriptionally, BAM cells showed strong immune activation, including enrichment of antigen presentation, leukocyte proliferation, and inflammatory pathways, with immune-associated changes also evident at the protein level at late stage (AD vs. MCI). In epithelial cells, transcriptional analyses revealed downregulation of cilia-related pathways, which was consistent with corresponding protein-level reductions in ADd. More broadly, transcriptional remodeling of barrier- and transport-associated programs—including tight junctions, extracellular matrix, cell adhesion, and ion/transport channels, was observed across fibroblast, endothelial, mural, and epithelial compartments, with these pathway classes likewise implicated in stage-associated proteomic remodeling. Together, these findings highlight three convergent molecular signatures of CP dysfunction during AD progression: BAM immune activation, broad remodeling of junctional, adhesion, ECM, and transport pathways, and cilia reduction.

### Inferred intercellular communication changes in the choroid plexus in AD dementia

To assess differences in intercellular signaling, we applied CellChat^24^ to snRNA-seq data to infer ligand–receptor–mediated communication. ADd cases showed higher weights of interactions compared with controls (Fig. 6a, number of interactions: Extended data Fig 5a). These increases were largely driven by differences in communication among fibroblasts, endothelial, immune, and mural cells through ECM-related pathways, including Collagen, Laminin, BMP, JAM, EPHA, and FN1 (Fig. 6b, Extended data Fig 5b-d, and Table 29).

**Fig. 6:**
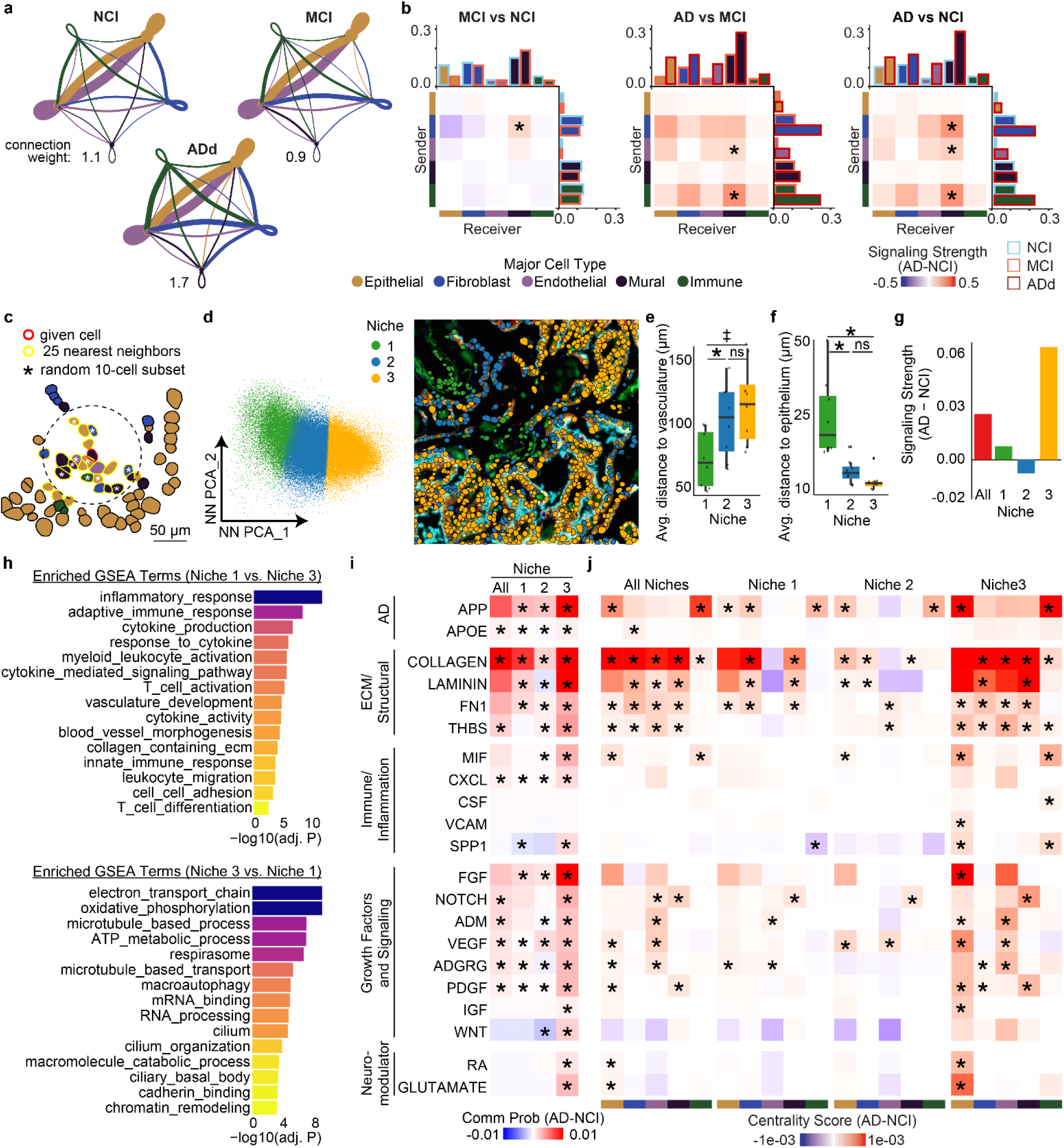
Altered CP intercellular communication in AD. **a,** AD differential CellChat inferred communication weights between major cell types throughout clinical disease progression in snRNA-seq. **b,** AD differential CellChat net probability (CellChat net$prob) heatmaps among all ligand-receptor pairs in each colored sender-receiver pair (* p < 0.05). **c**, Partial field of view (FOV #1.89), with cells colored by major cell type, to illustrate the nearest neighbor analysis approach. For each cell (e.g. red-outlined cell), a neighborhood transcriptome profile is calculated by averaging expression of a random 10-cell subset (white asterisks) of its 25 nearest neighbors (yellow-outline). **d**, Left: UMAP representation of neighborhood expression matrix colored by cluster (k-means = 3). Right: spatial plot of FOV #1.89, with cells colored by niche identity overlaid on immunofluorescent image. **e-f**, Boxplots of average cell to cell distances per niche to the nearest target cell type(s): Endothelial or Mural cell for (e), Epithelial cell for (f). Pairwise Wilcoxon tests with BH correction between niche (* p < 0.05, ‡ p=0.07, ns: not significant). **g**, CellChat normalized AD-differential signaling strength bar charts. **h**, Bar charts of select GO terms with positive enrichment (via fgsea on ranked t statistic) for niche 1 (upper) and niche 3 (lower). **i**, Interaction probability heatmap of select AD-differential CellChat pathways, for either the full or niche-specific datasets (* p < 0.05). **j**, For the same pathways in (i), AD-differential signaling split by colored cell types, for either the full or niche-specific datasets (* p < 0.05)

Because the CP is organized into stromal, vascular, and epithelial compartments, spatial context is essential for interpreting AD-related differences. We therefore performed nearest-neighbor analysis on spatial transcriptomics data to define local microenvironments (“niches”). Each neighborhood matrix was constructed by averaging expression across a random 10-cell subset of the 25 nearest neighbors, which revealed three spatial niches (Fig. 6c-d, Extended Data Fig. 6a-d): niche 1 corresponded to vessel-rich stroma (fibroblasts, endothelial, mural cells), niche 3 was enriched for epithelial cells, and niche 2 represented an intermediate epithelial environment (Extended Data Fig. 6e). Spatial mapping confirmed this organization, with niche 1 localized near vessels and niche 3 closer to the epithelium, which also showed the highest expression of *TTR* (Fig. 6e-f; Extended Data Fig. 6f). Examination of subtype distributions across niches revealed niche-specific enrichment of certain immune, fibroblast, and epithelial subtypes (Extended Data Fig. 6g). Gene set enrichment reflected these differences: niche 1 was enriched for immune activation, inflammation, vasculature, and ECM remodeling, while niche 3 was associated with metabolism, oxidative stress, autophagy, transport, and cilia (Fig. 6h).

We next applied CellChat analysis to the spatial transcriptomic data to resolve AD-associated signaling alterations across all cells and within defined spatial niches. Consistent with single-nucleus findings, overall cell–cell communication probabilities were higher in AD, particularly within niche 3 (Fig. 6g; Extended Data Fig. 6h). Across niches, the most upregulated pathways in AD involved APP-, APOE-, and ECM-related signaling (e.g., COLLAGEN, LAMININ, FN1, THBS), immune/inflammatory (MIF, CXCL), as well as growth factor–mediated networks (FGF, VEGF, PDGF, ADGRG), indicating widespread matrix remodeling and paracrine activation (Fig. 6i; Extended data Table 29). At the cell-type level, niche 1 and niche 2 showed more limited pathway activation, with significantly elevated pathways generally involving one or two dominant cell types (Fig. 6j). In contrast, niche 3 exhibited broader pathway activation that frequently involved multiple cell types (Fig. 6j), highlighting its role as a multicellular signaling hub in AD.

## Discussion

By integrating single-nucleus RNA-seq, ATAC-seq and spatial transcriptomic data, we establish a multi-omic atlas for a previously understudied brain border tissue—the human choroid plexus (CP). We defined 17 distinct CP cell states and revealed their molecular remodeling across the disease continuum from normal cognition to AD dementia. Together with proteomic datasets from CP tissue and cerebrospinal fluid, these results provide a publicly accessible, multidimensional resource for investigating CP biology, barrier regulation, and disease-associated remodeling in AD and human aging. Our multi-omic analyses revealed that AD dementia was not characterized by major differences in cell-type composition but by widespread cell-intrinsic transitions that collectively reshaped the CP’s immune, barrier, and stromal networks. These transitions converged into three major phenotypic axes —immune activation, barrier disruption, and stromal remodeling—that together define a chronically stressed and progressively compromised CP landscape in AD.

Along the inflammatory axis, both epithelial cells and border-associated macrophages (BAMs) exhibited progressive inflammatory activation and upregulated T-cell-activation, antigen-presentation, and cytokine-response programs, indicating coordinated immune engagement within the CP. At the late stage, BAMs apparently acquired stress-associated states marked by heat-shock-gene expression, suggesting maladaptive persistence under chronic inflammatory pressure. Consistent with these transcriptomic changes, immunostaining revealed increased CD68⁺ and HLA-DR⁺ macrophages indicating enhanced local macrophage activation. Despite macrophage upregulation of leukocyte-adhesion and antigen-presentation genes, we observed no change in CD3⁺ T cells—consistent with a sustained but compartmentalized inflammatory response distinct from the leukocyte trafficking phenotype characteristic of acute injury or infection^7,24,25^.

At the barrier level, our analyses revealed broad compromise of cell-junction and adhesion pathways across epithelial, endothelial, and fibroblast compartments. Epithelial cells showed selective downregulation of the tight-junction gene *CLDN5*, while pathway analyses indicated weakening of junctional integrity in secretory epithelial subtypes and endothelial cells, along with a reduction of the Mural_3 pericyte subtype, suggesting progressive loss of vascular support. In addition, fibroblasts—including the putative base-barrier Fibro_2 population—displayed lower expression of junctional components, supporting their contribution to barrier maintenance at the stromal–CSF interface. These multicellular alterations were accompanied by aberrant transporter and secretory pathways in epithelial cells along with a marked reduction in cilia function, validated by CROCC immunostaining, indicating impaired mechanosensory feedback control. Because motile and primary cilia help maintain epithelial polarity and barrier regulation through CSF sensing, their loss likely contributes to barrier dysfunction in AD. Consistent with these molecular and structural changes, CSF proteomic profiles revealed elevated inflammatory proteins together with reduced metabolic and ciliary-associated components, reflecting disrupted epithelial function and altered CSF composition in AD.

Finally, the remodeling axis highlighted extracellular-matrix (ECM) dynamics, reflecting both matrix degradation and restrained fibrotic deposition. Fibroblasts in AD displayed mixed transcriptional signatures of cell–matrix adhesion and collagen-containing ECM organization. Movat staining showed only a mild increase in collagen accumulation, contrasting sharply with the fibrotic scarring and persistent collagen deposition typical of acute injury or infection^26–28^. Other barrier-associated compartments—including epithelial, endothelial, and mural cells—also exhibited aberrant cell–matrix adhesion and extracellular-matrix pathways (Fig. 5j; Fig.6i), consistent with a coordinated structural remodeling rather than fibroblast-specific collagen deposition. These molecular alterations were further supported by proteomic analyses of CP tissue, which revealed parallel changes in ECM-related and adhesion proteins (Fig. 5j). Thus, the CP in AD exhibited a chronic, low-grade remodeling phenotype consistent with a long-term imbalance between matrix synthesis and degradation.

Taken together, these axes define a unified model of CP dysfunction in Alzheimer’s disease, characterized by chronic immune activation, barrier instability with CSF dysregulation, coordinated ECM and tissue remodeling. The coordinated transcriptional changes across epithelial, stromal, vascular, and immune compartments establish a foundation for mechanistic studies linking cilia function, chronic inflammation, and ECM turnover to disrupted CSF homeostasis. Collectively, these findings position the choroid plexus as a dynamic immune–barrier interface in AD, and highlight epithelial cilia maintenance, junctional resilience, and macrophage stress adaptation as conceptual entry points for therapeutic exploration.

## Acknowledgments

We are grateful to those who agreed to donate their brains for research. We thank all the employees at RADC for their support and assistance. This study was supported by NIA grants R01AG074082 and R01AG079223 (to Y.W.) and P30AG10161, P30AG72975, R01AG015819 R01AG017917, and U01AG61356 (to D. A. B.)

## Authors’ contributions

D.R.A., N.A.K., and S.D.T. designed and optimized snRNA-seq and snATAC-seq. D.R.A. and S.D.T. performed sample preparation and snRNA-seq and snATAC-seq experiments. S.D.T., H.V., and D.R.A conducted spatial transcriptomics (ST) experiments. J.X. performed snRNA-seq and ATAC-seq data pre-processing. T.J.P. performed all snRNA-seq and snATAC-seq data analyses and generated related figures. T.J.P. Performed ST data preprocessing. T.J.P. and D.R.A. conducted ST analyses and generated related figures. N.A.K. contributed to snRNA-seq analyses. H.V., D.S., A.K.L., and N.A.K. performed immunohistochemistry experiments and data analysis. D.D. and N.T.S. generated mass spectrometry proteomics data. B.N. conducted proteomics analysis. T.J.P., D.R.A., and Y.W. wrote the manuscript with input from all co-authors. D.A.B. directed the parent ROSMAP study. Y.W. conceived and supervised the overall project.

## Competing interests

The authors declare no competing interests.

## Corresponding authors

Correspondence to Yanling Wang

## Data availability

Sample information, cluster maker genes, and differentially expressed genes are provided in the Supplemental Information. All snRNA-seq, snATAC-seq and spatial transcriptomic data are available at Synapse under accession code syn2580853. ROS/MAP data can be requested by qualified investigators (www.radc.rush.edu).

## Code availability

Code for analysis is available at GitHub (https://github.com/Tristan-J-Philippe/Multi-omic_Choroid-Plexus_Alzheimers-Disease).

## Online Methods

### Tissue collection

Human post-mortem choroid plexus (CP) tissue was collected from the lateral ventricles of 69 participants in the Religious Order Study Rush Memory and Aging Project (ROSMAP)^16^. The tissue was bisected and preserved for snRNA-seq, proteomics, IHC, or spatial transcriptomics. Tissue used for snRNA-seq and proteomics was snap frozen and stored at −80°C whereas tissue for immunostaining and spatial transcriptomics was formalin fixed and paraffin embedded. Clinical diagnostic, pathologic AD and cerebral amyloid angiopathy (CCA) were assessed as previously described^16,29–31^. The Institutional Review Board of Rush University Medical Center approved both studies. All participants signed an informed consent and Anatomic Gift Act.

### Nuclei isolation for snRNA-seq

The clinical diagnosis was used to group ADd, MCI, and NCI into batches of 4-6 participants for snRNA-seq (n=69; Fig.1b, Extended Data Table 1); these groups were balanced for age and sex. All procedures were carried out on ice or at 4 °C to minimize RNA degradation, unless otherwise specified. 100 mg frozen CP tissue was minced on dry ice, then thawed in 0.1X lysis buffer (250 mM sucrose, 25 mM KCl, 5 mM MgCl2, 10 mM Tris pH8, 1μM DTT, 15μM actinomycin, 0.2U/μL RNAse inhibitor, 0.01% v/v Triton X-100). Tissue was pressed through a 100μm cell strainer with the back end of a 5mL plunger, rinsing every 30 seconds with 200 μL fresh 0.1X lysis buffer for 3 min total per sample. The cell suspension was pelleted at 600g for 4 min (all centrifugation steps were at 4°C), supernatant was removed, and pellets were resuspended in 1.5 mL 1X lysis buffer with 0.1% Triton X-100. Nuclei were homogenized using a Dounce homogenizer (10 times using pestle B) and incubated for 2 min on ice. Nuclei were washed twice: centrifuged at 700g for 4 min and resuspended in 1 mL of wash buffer (1.5% BSA, 0.2U/μL RNAse inhibitor in DPBS). Samples were purified by iodixanol gradient (37%, 29%, 25% with sample) centrifugation (13500g, 20min, OptiPrep Sigma) and collected from the 29-37% interface, washed once, filtered through a 40 μm cell strainer (Flowmi), and washed again before QC and counting using Trypan blue.

For barcoding, nuclei (100k in 187.5uL total volume * 2 unique barcodes per sample) were incubated with 4 μM of anchor and 4 μM of a unique barcoded oligonucleotide and incubated for 5min (final concentration of 50 nM of each; see Nuclei hashing design). Followed by 4 μM co-anchor, mixed by tapping, incubated on ice for 5 min, and 3 washes. Hashed nuclei were then combined, diluted to ∼40k nuclei in 43.2 μL, and super-loaded (∼40k nuclei loaded per lane) onto 4 lanes of 10X Genomics Chip G or Chip J. Libraries were prepared using the 10x Genomics 3’ single-cell gene expression assay v3.1 (CG000317 Rev C) for snRNA-seq alone and (CG000338) for single nuclei RNA and ATAC sequencing except as detailed in method: Nuclei hashtag, followed by sequencing on a NovaSeq 6000 to an average read depth of 50k RNA reads/cell and 15k hash reads/cell.

### Nuclei hashtag oligonucleotide design and library preparation

Nuclei were barcoded using a modified MULTI-seq^32^ protocol to enable pooling and super-loading of the 10X chip. The Anchor sequence was: /5Chol-TEG/GTAACGATCCAGCTGTCACTAGATCGGAAGAGCACACGTCTG, where ‘5Chol-TEG’ denotes a 5’

Cholesterol modification conjugated via a triethylene glycol linker. The Co-Anchor sequence was: AGTGACAGCTGGATCGTTAC/3CholTEG/. Sixteen barcoded oligonucleotides (aka “hashtag oligos”) were designed following the pattern R2-BC-poly(A), where R2 was a constant partial Read 2 sequence (CAGACGTGTGCTCTTCCGATCT), BC was a unique sample-specific 15-nt barcode sequence (sequences listed below), and poly(A) was a 30-nt poly(A) sequence. The poly(A) enables recovery of hashtag oligos along with mRNA following reverse transcription. To further enrich for hashtag oligos, a Truseq 2 primer (CAGACGTGTGCTCTTCCGATC) was spiked into the cDNA amplification reaction (CG000317 Rev C step 2.2) at 100 nM. All oligonucleotides were ordered from IDT. For the hashtag library sample index PCR (step 3.5), the reaction consisted of 50 µl Amp mix, 40 µl EB buffer, 5 µl cDNA from transferred supernatant cleanup (step 2.3B), and 5 µl dual index from plate TT. PCR was run for 9 cycles with default settings, followed by a 1.2X SPRI cleanup. Next, 20 ng (∼1 µl) of PCR product was used for an additional sample index PCR (6 cycles) to enrich for full-length product. The reaction also consisted of 30 µl Amp mix, 1.2 µl each of Illumina P5 and P7 primers (10 µM), and EB buffer to a total volume of 60 µl. This was followed by one 0.9X SPRI cleanup to remove free primer and primer dimer. The hashtag (HT) libraries (∼242-bp total size) were mixed with GEX libraries (ensuring distinct TT indices) at a molar ratio of 1:6 HT:GEX in the final pool to achieve the desired ∼1:3 ratio of sequencing reads.

Sample barcodes:

AAGGCAGACGGTGCA

GGCTGCGCACCGCCT

GACGCGCGTTGTCAT

GAAGAAGCGTTATTC

AGAGCTAGGATCGGA

GCTAGCGTAATGTGG

CGGTTAGCGCGGTTC

AGGCGTGCACGTGGT

CACATATAGAATTAG

ATAGCGGCTCTATCG

AGCGCATAGTTATCT

CAGGTCGGCCAAGAT

GCCAAGATCAGGTCC

TAGGTTACGAGTGTG

ATAATCATTACGTGG

ATCGAACCGACAGAG

### Spatial transcriptomics of CP using Nanostring CosMx

Eight human CP samples were selected (4 NCI, 4 AD), matched for sex, and balanced for age. 5-μm thick FFPE sections were mounted on Leica BOND Plus microscope slides (4 sections per slide; 2 slides total), ensuring that the tissue resided within the 15x20mm imaging area. Slide preparation followed the manufacturer’s protocol (Nanostring MAN10159-03) with the following modifications. Antigen retrieval was carried out using a steamer for 15 min once the retrieval solution reached 99°C. Tissue permeabilization was performed for 15 min at 40°C using 3 μg/mL Proteinase K. Probe hybridization occurred for 18 hr at 37°C.

Primary antibodies included Cell Segmentation Mix 1 (CD298/B2M), Marker Mix 1 (PanCK/CD45), and Marker Mix 2 (CD68) at the recommended dilutions and were incubated for 90 min (instead of 60 min) at room temperature. To load slides on the Nanostring CosMx instrument, we followed the Instrument User Manual (MAN-10161-03-2; software version 1.2) using configuration B for pre-bleaching and configuration A for cell segmentation. Field of view (FOV) selection encompassed the entire tissue-covered area.

### TMT-based proteomics of choroid plexus tissue

A subset of choroid plexus samples (n = 16; Extended Data Table 1), balanced for age and sex, were analyzed by mass spectrometry. Briefly, tissues were lysed in 8 M urea, 100 mM Tris, 100 mM NaH₂PO₄ (pH 8.5) with protease and phosphatase inhibitors (Thermo Fisher), reduced with 5 mM dithiothreitol, alkylated with 10 mM iodoacetamide, and digested overnight with trypsin. Peptides were labeled using the TMTpro 16-plex kit (Thermo Fisher Scientific), pooled, and fractionated by high-pH reversed-phase chromatography. Fractions were analyzed on an Orbitrap Lumos mass spectrometer (Thermo Fisher Scientific) coupled to a 3000 RSLC nano-UPLC system using a 26-min gradient on a 1.7 µm C18 column (Waters). MS scans were acquired at 120,000 resolution with a 3 s cycle time, and the top precursors (charge 2–5) were selected for MS/MS with higher-energy collisional dissociation (HCD 35%), 50,000 resolution, 125k AGC target, and 86 ms maximum injection time, with dynamic exclusion of 20 s. Raw files were processed in Proteome Discoverer v3.0 using Sequest HT against the UniProt human proteome database for TMT-based quantification, and protein abundances were normalized across channels for downstream analysis.

### Olink-based proteomics of cerebrospinal fluid

Cerebrospinal fluid (CSF) samples (n=24; Extended Data Table 1) were analyzed using the Olink Explore 384 platform (Thermo Fisher Scientific). The following panels were included: Cardiometabolic, Cardiometabolic II, Inflammation, Inflammation II, Neurology, Neurology II, Oncology, and Oncology II. Data processing, QC, and normalization were performed using the Explore Calculation Module v3.8.1, applying intensity normalization on a log₂ scale.

### snRNA-seq

We ran bcl2fastq, followed by CellRanger v7.0.1 to generate RNA, ATAC, and hash count matrices using the GRCh38 genome. Downstream analyses were conducted in R Seurat v4.2.0 package^33^. RNA and ATAC count matrices were treated separately to retain nuclei with sufficient modality-specific quality. For snRNA-seq we excluded droplets with >12,000 or <500 features and <500 counts. Ambient RNA contamination was corrected using SoupX v1.6.2^34^ with autoEstCont (tfidMin=0.7–0.9) to ensure at least 100 genes were used for estimation. Nuclei were demultiplexed using HTOdemux, which also flagged hash doublets.

In alignment with previously established strategies for guided doublet removal^35,36^ we leveraged previously reported markers for the choroid plexus, brain vasculature, and brain immune cells in mouse and human (see Extended Data Table 30). Data were normalized and scaled incorporating the top 3k variable features and reported markers, which were then used for Principal Component Analysis (PCA, top 30 embeddings), Harmony^37^ v0.1.0 integration by sequencing library, dimensionality reduction (UMAP), and clustering (Louvain algorithm, res=0.03-0.7)^38^. Positive marker genes were identified using a bimodal Wilcox rank sum-test (log_2_FC>0.25, p<0.05). Since ‘dark’ epithelial cells are characterized by high mitochondrial content^39,40^, mitochondrial content >5% were only retained in the epithelial cluster. Nuclei with mixed cluster signatures were excluded, known doublets were removed and misclustered cells were removed from UMAP plots. Data were iteratively reprocessed as above but in an unsupervised manner with the top 3k unknown variable features to identify major and sub cell types.

### snATAC-seq

Following the generation of raw count matrices, we utilized Seurat v4.2.0^33^ and Signac 1.10.0^41^ . Low-quality droplets with total fragments <100 or >5,000, blacklist ratio >0.1, nucleosome signal >4, and transcription start site (TSS) enrichment score <3 was excluded. After demultiplexing with HTODemux, doublets were removed using the semi-supervised method described above (Methods: snRNA-seq). Chromatin accessibility data were normalized using RunTFIDF, reduced with RunSVD, and the top 30 components were used for dimensionality reduction and clustering. To annotate clusters, we generated a pseudo-expression “activity” matrix, which was normalized, scaled and a bimodal Wilcoxon rank-sum test (log₂FC > 0.25, p<0.05) was used to identify cell type markers. We annotated chromatin features with the UCSC Genome Browser, motif analysis with motifmatchR v1.31.1^42^ (JASPAR 2022, default thresholds), and transcription factor activity with Signac Footprint (default settings).

### Spatial transcriptomics

Initial preprocessing followed the AtoMx pipeline (MAN-10162-03; software v1.2). Cellpose v2 was used for segmentation (non-neuro human tissue, cell dilation: 2.17 μm, cell diameter: 7.28 μm, nucleus size: 7.2 μm, membrane size: 5.7μm, Models: cyto2, flow threshold: 0.1, probability threshold: 2, min. size: 3.6μm) followed by QC and flat file export to Seurat v5.1.0^43^. Segmented cells with <100 counts, >3500 counts, <50 features, >2000 features, qcCellsFlagged==T, or an area <10μm^2^ were excluded. We used SCTransform v.0.4.1^44,45^ to correct, scale, and normalize; and harmony to correct for individual and slide effects. The semi-supervised method described above (Methods: snRNA-seq) was used to identify potential segmentation errors (e.g. cells with high expression of marker genes from multiple major cell types). These were excluded for downstream cell type annotation (unsupervised, top 2k) and AD-differential testing but retained for spatial neighborhood and CellChat analysis. Due to limitations of unsupervised clustering algorithms in resolving rare cell types (<6,000 cells per major cell type), we applied a supervised mapping approach using the snRNA-seq dataset as reference. Specifically, spatial transcriptomics data were annotated using RCTD^23^ with default parameters, leveraging pseudobulked snRNA-seq expression profiles for each major cell type as reference.

### Differential expression analyses

We performed differential gene expression, protein abundance, and chromatin accessibility analyses using a linear modeling framework with empirical Bayes moderation as implemented in voom and limma v3.54^46^. Where applicable, models were adjusted for age, sex, postmortem interval (PMI), and sequencing library. Genes and proteins with fewer than two counts per sample or expressed in fewer than 1% of cells were excluded. Accessible chromatin regions and gene activity scores with variance-standardized transformation (vst) scores below 1 were removed. For the snRNA-seq dataset, samples with >100,000 total pseudobulked counts were split by sequencing library to create technical replicates; samples with <60,000 counts were excluded. Genes with a false discovery rate (FDR) <0.01 and absolute log2 fold change (LFC) >0.5 were considered differentially expressed. For smaller datasets (spatial transcriptomics, chromatin accessibility (snATAC-seq), and proteomics) validation thresholds were relaxed (p < 0.05, LFC > 0.2). We tested cell type specific transcriptional signatures and correlated these between modalities. We tested contrasts between clinical diagnostic (cogdx), neuropathological categories (adnc_ad), and cognitive decline (last cognitive score before death) among major and minor cell types.

### Enrichment analysis

Ranked gene lists were stratified by directionality and tested using fgsea v1.25^47^ against Gene Ontology (GO) Biological Process, Molecular Function, and Cellular Component pathways. A normalized enrichment score (NES) >0.6 and FDR <0.15 for spatial transcriptomics, snATAC-seq, and proteomics were used to define significant pathways. Enrichment and differential expression results were visualized using ClueGO v2.5.10^48^, Cytoscape v3.10.2, ComplexHeatmap^49,50^, Seurat, and ggplot2 v3.4.4^51^.

### Proportion analysis

As previously described^52^, we applied a linear modeling with empirical Bayes moderation to assess cell type proportions as a function of cognitive diagnosis (cogdx). Where applicable, models were adjusted for parenchymal contamination, age, sex, and PMI. Statistical significance was defined as FDR < 0.01.

### Pattern matching in snRNA-seq

We applied a template matching algorithm^22^ to identify genes associated with disease progression along three possible positive and negative trajectories. Differentially expressed genes (p<0.05, LFC>0.25) from the MCI vs. NCI, ADd vs. MCI, and ADd vs. NCI contrasts were classified as early-stage (significant in MCI vs. NCI and ADd vs. MCI, but not AD vs. NCI), late-stage (significant in ADd vs. MCI and ADd vs. NCI, but not MCI vs. NCI), and progressive (significant in MCI vs. NCI and ADd vs. NCI) patterns. Genes were correlated with the corresponding stage template, and only those with p < 0.05 were retained.

### Nearest Neighbor analysis

To obtain a better understanding of cell-cell interaction we included cells with mixed signatures and mapped cells to their closest major cell type (Methods: Spatial transcriptomics data processing), only excluding cells that did not meet QC metrics, parenchymal cells and red blood cells. This resulted in 166,607 total cells for analysis. A spatial neighborhood matrix was generated using CellularNeighborhoods v0.0.0.9000 available here: https://github.com/Nanostring-Biostats/CosMx-Analysis-Scratch-Space/tree/Main/_code/cellular-neighborhoods. Each cell’s neighborhood expression was calculated as the mean expression of a random 10-cell subset of its 25 nearest neighbors, based on cell centroid x-y coordinates. We performed normalization/scaling, PCA (50 components), Harmony (50 components, accounting for slide batch, individual, and AD status), UMAP, and clustering (kmeans = 3) to obtain 3 spatial neighborhoods or ‘niches’.

### CellChat

We used CellChat v1.6.1^24^ to infer intercellular communication networks among major cell types stratified per cognitive diagnosis (MCI, MCI, and ADd) with default parameters for the snRNA-seq dataset and for spatial transcriptomics after nearest neighbor analysis. For spatial transcriptomics, CellChat was run separately for each niche with SCTransformed counts, computeCommunProb with a contact.range=20 microns, interaction.range=250 microns, scale.distance=0.69 (empirically determined), and all other parameters default. We also normalized cellchat metrics (interaction weight, net$prob, netP$prob, and centrality score) by the number of cell pairs with the potential for interaction (e.g. within 250 microns), thereby accounting for differences in cell number and density between NCI and AD, as well as across niches.

To identify disease-associated signaling pathways, objects for each cognitive diagnosis were merged and comparative network analyses performed. A customrankNet() function was implemented to rank pathways with p<0.05 by summed communication probability (weight) per condition, calculating relative contributions across diagnostic groups. Pathway-level metrics (information flow and interaction counts) were extracted from the netP slot of the merged object. To assess cell-type-specific signaling changes, centrality scores (incoming + outgoing) were computed for each cell type and condition using netAnalysis_computeCentrality(). To ensure robust conclusions, previously identified (Methods: Differential analyses: Expression) statistics for each ligand- receptor pair were utilized and the average log_2_FC of ligand and receptor(s) was calculated. We removed non-directional or low-confidence interactions (ligand-receptor pairs with absolute combined log₂FC < 0.05, ligand detected in < 5% of sender cells, receptor detected in < 5% of receiver cells, and/or net$p > 0.05). At the pathway level, median log_2_FC was calculated per pathway and a Wilcoxon signed-ranked test was applied to pathways with ≥ 3 interactions followed by FDR correction. Pathways defined as significant had absolute median log_2_FC ≥ 0.15, FDR ≤ 0.01, absolute AD-differential net centrality > 0.0001, and matched centrality and median log_2_FC directionality.

### Cell-cell distance-based analysis

We converted centroid x-y coordinates from pixels to microns (x0.18) and computed per-cell distances to the nearest cell type (e.g. Mural/Endothelial, Epithelial, Parenchymal), then calculated and plotted per-individual averages across spatial niches or cell subtypes. We performed a Kruskal-Wallis test and used Dunn’s test with Benjamini-Hochberg correction for multiple testing.

### Immunohistochemistry

Human FFPE CP sections were stained using a Bond-Rx research stainer (Leica Biosystems). Briefly, slides were baked for 60 min at 60°C prior to dewaxing for 30 s at 72°C. For multiplexed cell typing, an Opal 6-plex detection kit (Akoya Biosciences #NEL871001KT) was used as follows; slides were incubated with CD45 (1:50) for 30 min followed by 10 min Opal polymer HRP Ms+Rb and 10 min Opal signal development with Opal 620 diluted at 1:150. Slides were antigen retrieved for 20 min in pH 6.0 citrate buffer solution (Leica Biosystems #AR9961) and the process was repeated with DCN (1:100; Opal 570 (1:150)), TTR (1:500; Opal 690 (1:150)), PECAM (1:50; Opal 520 (1:150)) and ACTA2 (1:1000; Opal 480 (1:150)). For ciliary rootlet structures, slides were subjected to HIER with a pH 9.0 EDTA buffer solution (Leica Biosystems #AR9640) for 20 min at 100°C followed by 10 min peroxide block. CROCC (1:100) and TTR (1:500) were incubated together for 60 minutes followed by 60 min of secondary anti-rabbit Alexa Fluor 647 (1:1000) for CROCC. The TTR antibody was detected with a Mouse IgG-HRP secondary, incubated for 60 min, followed by a 5 min incubation with TSA Fluorescein. All slides were then incubated with Trueblack quencher for 1 minute, counter stained with spectral DAPI for 5 min, and mounted. Imaging was performed using a PhenoImager (Akoya Biosciences) at 40X. ROIs were selected using Phenochart 2.2.0 (Akoya Biosciences) and spectrally unmixed composite TIFs were generated using inForm (Akoya Biosciences).

For CROCC puncta analysis, images were imported into ImageJ and split into individual channels. CROCC channel image was converted into 8-bit and threshold was set to mask only the puncta. Analyze particles tool in ImageJ was run with the following parameters: Size=1.5-25.0 μm and Circularity= 0.1-1.0. The results were exported and analyzed in GraphPad prism (v10.5.0). For classification of TTR+ epithelial cells, the raw composite TIFs were analyzed using Qupath (v0.5). Cells were defined in each ROI using QuPath Cell Detection tool with the following parameters, requested pixel size = 0.5μm, background radius = 8μm, median filter radius =0μm, Sigma value for Gaussian filter = 1.5μm, minimum nucleus area = 10μm^2^, maximum nucleus area =400μm^2^, and intensity threshold = 2. Cell parameters include a cell expansion of 5μm from the cell nucleus, and general parameters include smoothing of the detected nucleus/cell boundaries. Shape, intensity and smoothing features were added for all detections, and the “Train object classifier” tool was used to classify the TTR+ epithelial cells. All measurements were exported and analyzed in GraphPad prism (v10.5.0).

### Multiplex immunohistochemical staining using Akoya Phenocycler fusion (PCF)

Human FFPE CP sections were stained using the Akoya Biosciences Phenocycler Fusion 2.0 workflow as per manufacturer’s instructions. Briefly, the slides were baked overnight at 60 degrees and de-waxed, hydrated and antigen retrieval performed in a pressure cooker for 20 min in AR9 buffer. Staining was performed for 3 hrs at room temperature followed by washing and post-fixation steps. Imaging was performed on Akoya Phenocycler Fusion 2.0 instrument at 20X. Qptiff image files generated were imported into QuPath (v0.5) for quantitative analysis.

### Phenocycler Fusion Image Analysis

For analysis of CD163+ BAM cells, ROIs were defined in each image and QuPath Cell Detection tool was used with following parameters: detection channel= CD163, requested pixel size = 0.5μm, background radius = 20μm, median filter radius =0μm, Sigma value for Gaussian filter = 1.5μm, minimum nucleus area = 15μm^2^, maximum nucleus area =400μm^2^, and intensity threshold = 50. No cell parameters were selected, and general parameters include smoothing of the detected nucleus/cell boundaries. HLA-DR and CD68 intensity measurements from CD163^+^ cells were exported and analyzed in GraphPad Prism (v10.5.0).

For T-cell analysis, ROIs were defined in each image and QuPath Cell Detection tool was used with following parameters: detection channel= CD3e, requested pixel size = 0.5μm, background radius = 20μm, median filter radius =0μm, Sigma value for Gaussian filter = 1.5μm, minimum nucleus area = 10μm^2^, maximum nucleus area =400μm^2^, and intensity threshold = 50. No cell parameters were selected, and general parameters include smoothing of the detected nucleus/cell boundaries. Shape, intensity and smoothing features were added for all detections, and the “Train object classifier” tool was used to classify the CD4+ and CD8+ T-cells. All T-cells were manually annotated based on whether they are present inside (non-infiltrating) or outside (infiltrating) of Caveolin/Collagen IV+ vessels. The numbers of infiltrated T cells were plotted relative to the area of the ROIs and analyzed in GraphPad Prism (v10.5.0).

### Movat’s Pentachrome staining and analysis

Human FFPE CP sections were stained with Movat’s Pentachrome stain kit (Abcam, ab245884) as per the manufacturer’s instructions in two batches. Slides were imaged using the Phenocycler Imaging system (Akoya Biosciences) at 40X. 4-10 ROIs (928μm x 697μm) were selected from each case using Akoya Phenochart 2.2.0 and RGB TIFs were exported using inForm (Akoya Biosciences).

Images were processed using the Colour Deconvolution Package (https://blog.bham.ac.uk/intellimic/g-landini-software/colour-deconvolution-2/) in ImageJ (v1.54) to extract the yellow channel. Images were converted to 8 bit and exported as image txt files for further processing in R. Background subtraction was performed to align batches, and pixels that did not correspond to tissue were excluded. For each ROI, the distribution of pixel intensities was calculated. The pixel distributions for each individual were averaged to present the distribution for each disease group, plotted with ggplot2. The AUC for each ROI at high collagen (cutoff of 200) was calculated using the trapz() function from the pracma library (v2.4.4) and a one-way ANOVA performed to test significance.

## Extended Data Figures

**Extended Data Fig. 1:**
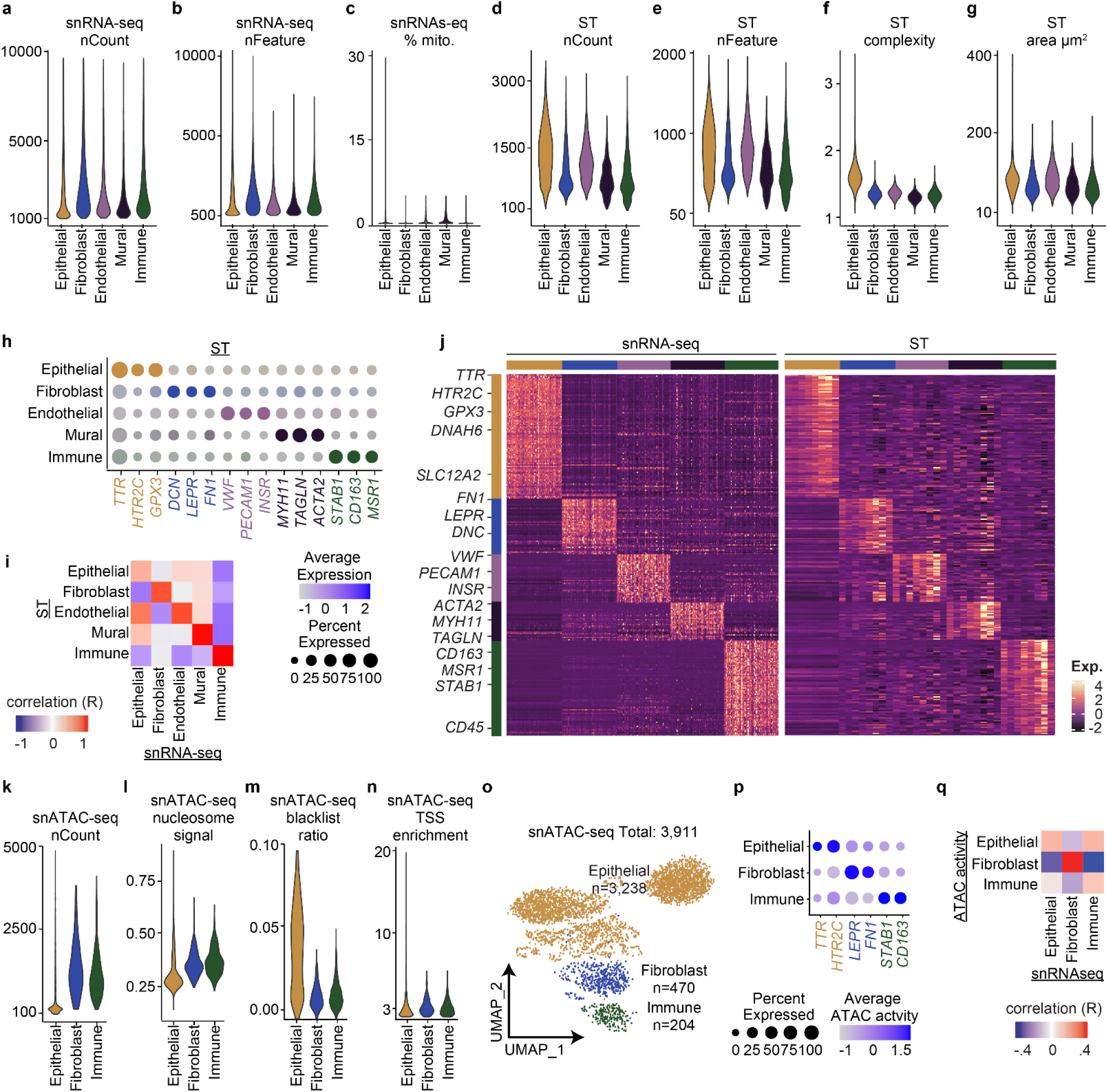
QC and cell-type validation across snRNA-seq, ST ad snATAC-seq. **a–c,** Total RNA counts(a), number of detected features (b), and Mitochondrial read percentage (c) violin plots per snRNA-seq subtype. **d–g,** Total RNA counts (a), number of detected features (b), segmented cell area (µm²; c) and complexity (d) violin plots per spatial transcriptomic (ST) subtype. **h,** Unsupervised spatial transcriptomic confirmation of canonical marker expression dot plot across cell clusters. See Extended Data Table 4. **i,** Pearson correlation of ST to RNA expression by major cell type.. **j,** Heatmap of pseudobulked marker gene expression (z-score) identified in snRNA-seq (limma; FDR < 0.01, log₂FC > 0.25). Heatmap of the same gene set in pseudobulked ST. **k-n,** Total peaks counts (k), nucleosome_signal (l), blacklist_ratio (m), and TSS_enrichment (n) violin plots per snATAC-seq subtype. **o,** UMAP projection of major celltypes with cell counts indicated. **p,** Unsupervised snATAC-seq confirmation of canonical marker expression dot plot across cell clusters. See Extended Data Table 6. **q,** Pearson correlation of chromatin accessibility to RNA expression pooled among major cell types.

**Extended Data Fig. 2:**
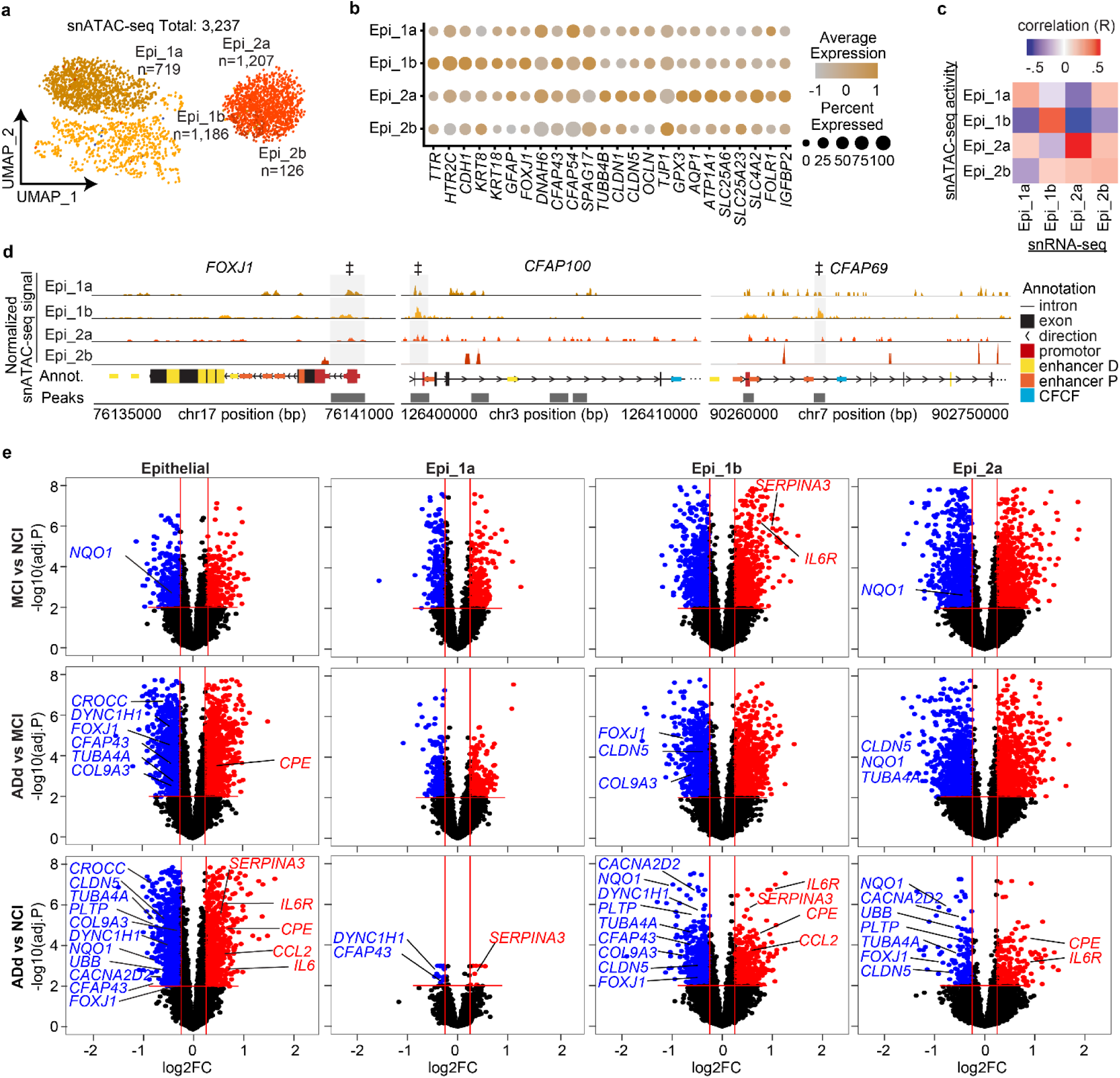
Epithelial cell subtypes defined by snATAC-seq and differential gene expression across disease stages **a,** snATAC-seq UMAP colored by epithelial subtype. **b,** Unsupervised snATAC-seq epithelial subtype-defining marker dot plot (FDR < 0.01, log₂FC > 0.25). See Extended Data Table 9. **c,** Pearson correlation of chromatin accessibility (ATAC activity) to RNA expression pooled among epithelial subtypes. **d,** Differentially accessible chromatin peaks for select cilia genes per epithelial subtypes (‡ p<0.05). Genome specific annotations and accessible peaks are shown below. **e,** snRNA-seq **e**pithelial and **s**ub type * disease specific DEG volcano plots. Select genes representing key GO pathways are labelled (all FDR < 0.01 and log₂ fold change > ±0.25). **f,** Scatter plots contrasting t-statistics from differential gene expression analysis between clinical stages (MCI vs NCI and ADd vs NCI), Select genes representing key GO pathways are labelled (all FDR < 0.01 and log₂ fold change > ±0.25).

**Extended Data Fig. 3:**
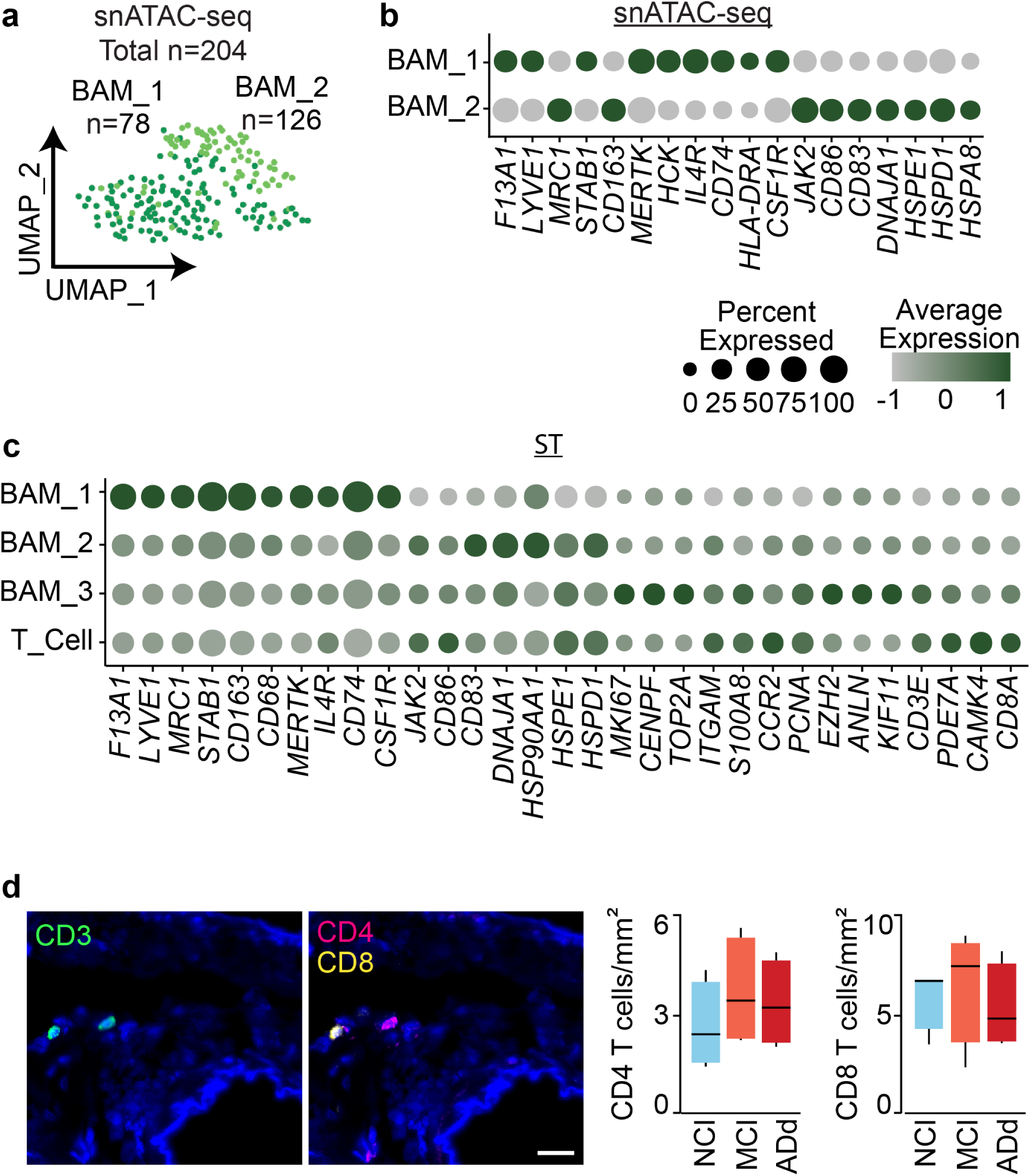
Immune cell subtypes defined by snATAC-seq and ST. **a,** snATAC-seq UMAP colored by immune subtypes. **b,** Unsupervised snATAC-seq immune subtype-defining marker dot plot. See Extended Data Table 16. **c,** Supervised spatial transcriptomics (ST) immune subtype-defining marker dot plot. **d,** CD4 and CD8 staining and quantifications across NCI, MCI and AD groups.

**Extended Data Fig. 4:**
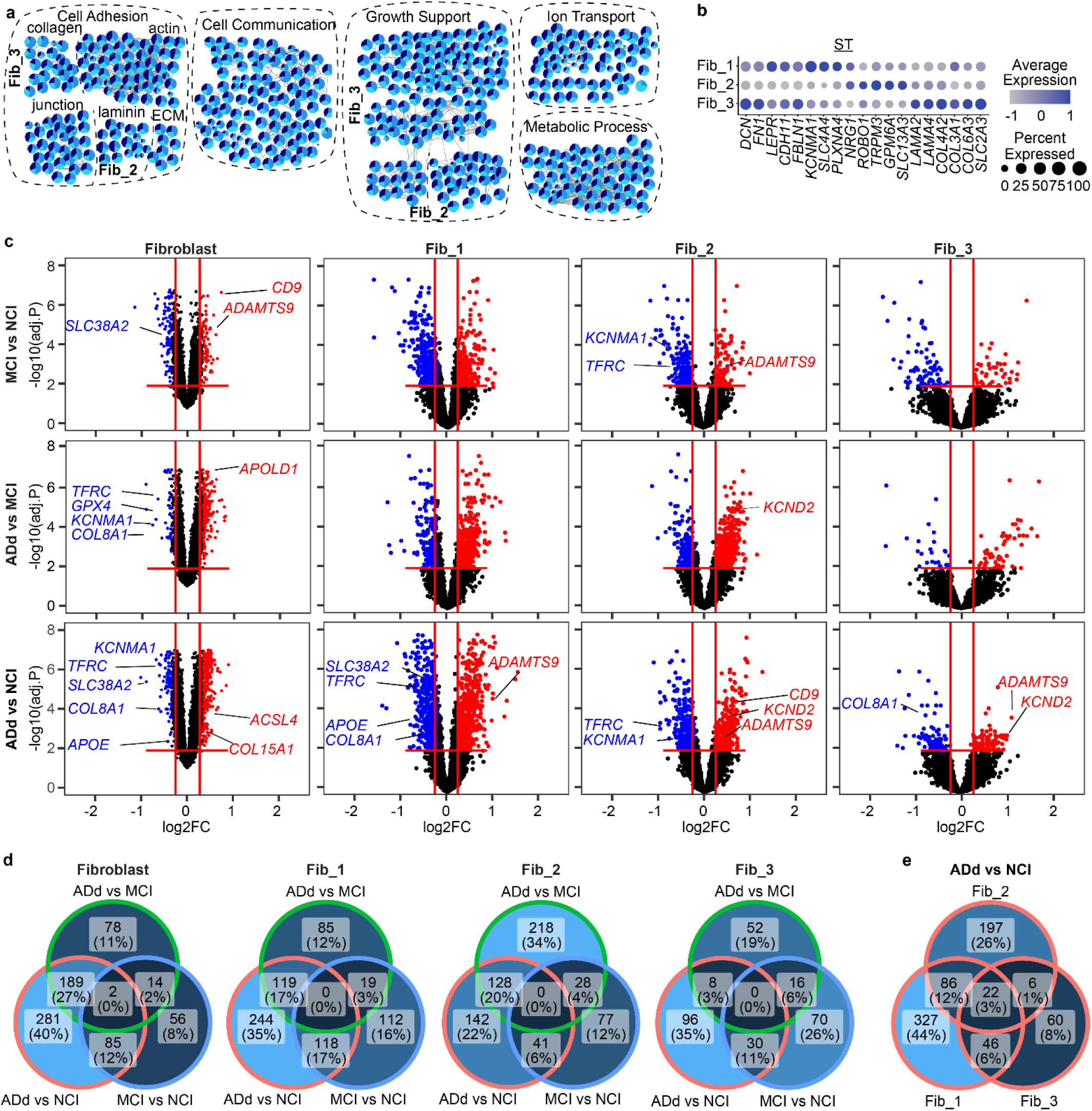
Fibroblast subtype-resolved differential gene expression across disease stages. **a,** Fibroblast subtype GO enrichment network plot (ClueGO on subtype-defining markers) grouped by functional category. **b,** Supervised spatial transcriptomics (ST) fibroblast subtype-defining marker dot plot. **c,** snRNA-seq fibroblast and **s**ub type * disease specific DEG volcano plots. Select genes representing key GO pathways are labelled (all FDR < 0.01 and log₂ fold change > ±0.25). **b,** Overlap of differentially expressed genes across subtypes and disease stages.

**Extended Data Fig. 5:**
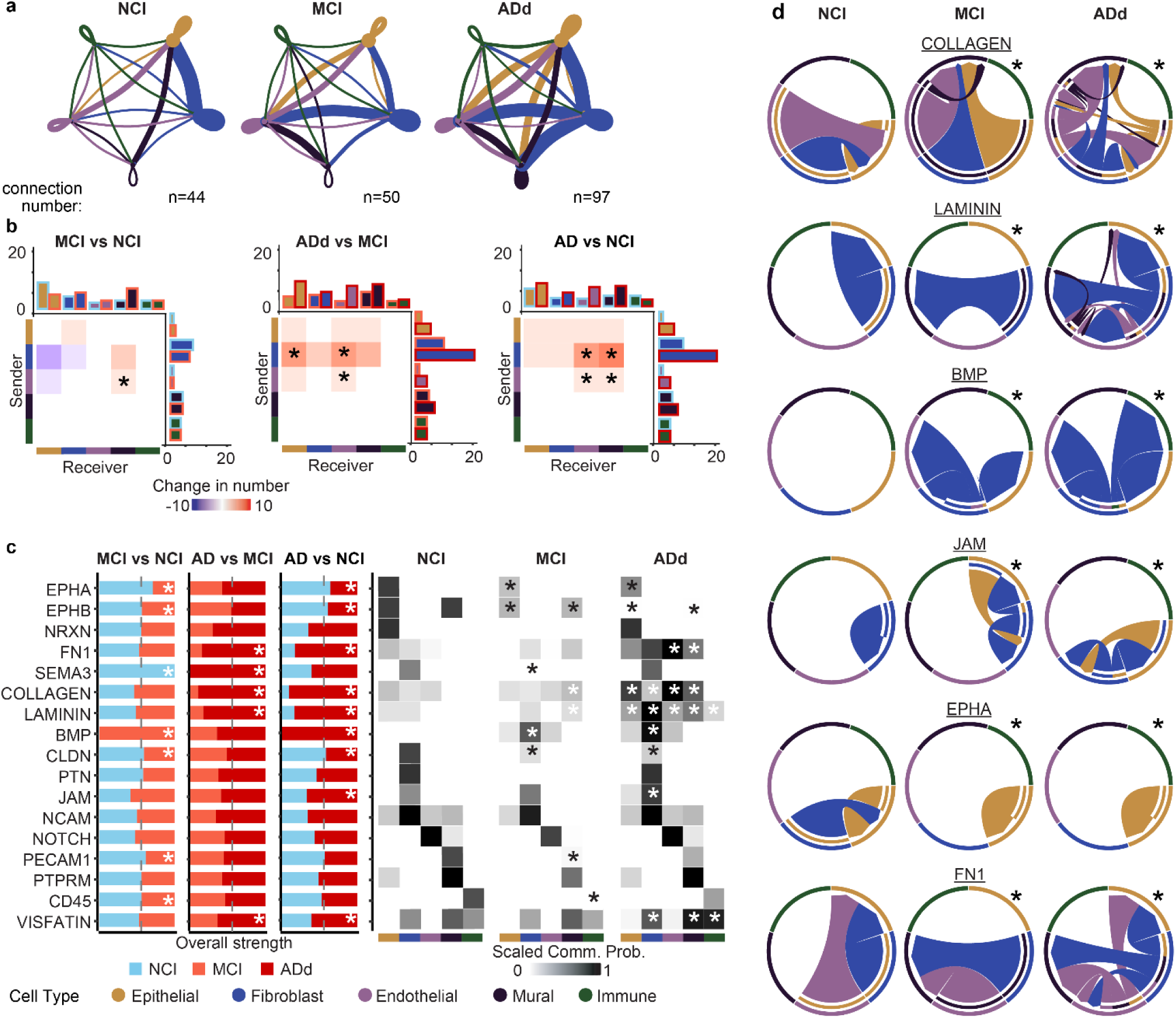
CellChat analysis of snRNA-seq data reveals altered CP intercellular communication in AD. **a,** Differential CellChat inferred communication number between major cell types throughout clinical disease progression in snRNA-seq. **b,** AD differential CellChat communication number heatmaps among all ligand-receptor pairs in each colored sender-receiver pair (* p < 0.05). **c,** Select signaling pathways mediating differential intercellular communication throughout clinical disease progression. Left: Total signaling strength histogram. Right: Scaled communication probability per cell type (* p < 0.05). See Extended Data Table 29. **d,** Selected chord diagrams of intercellular signaling between major cell types throughout clinical disease progression (* p < 0.05).

**Extended Data Fig. 6:**
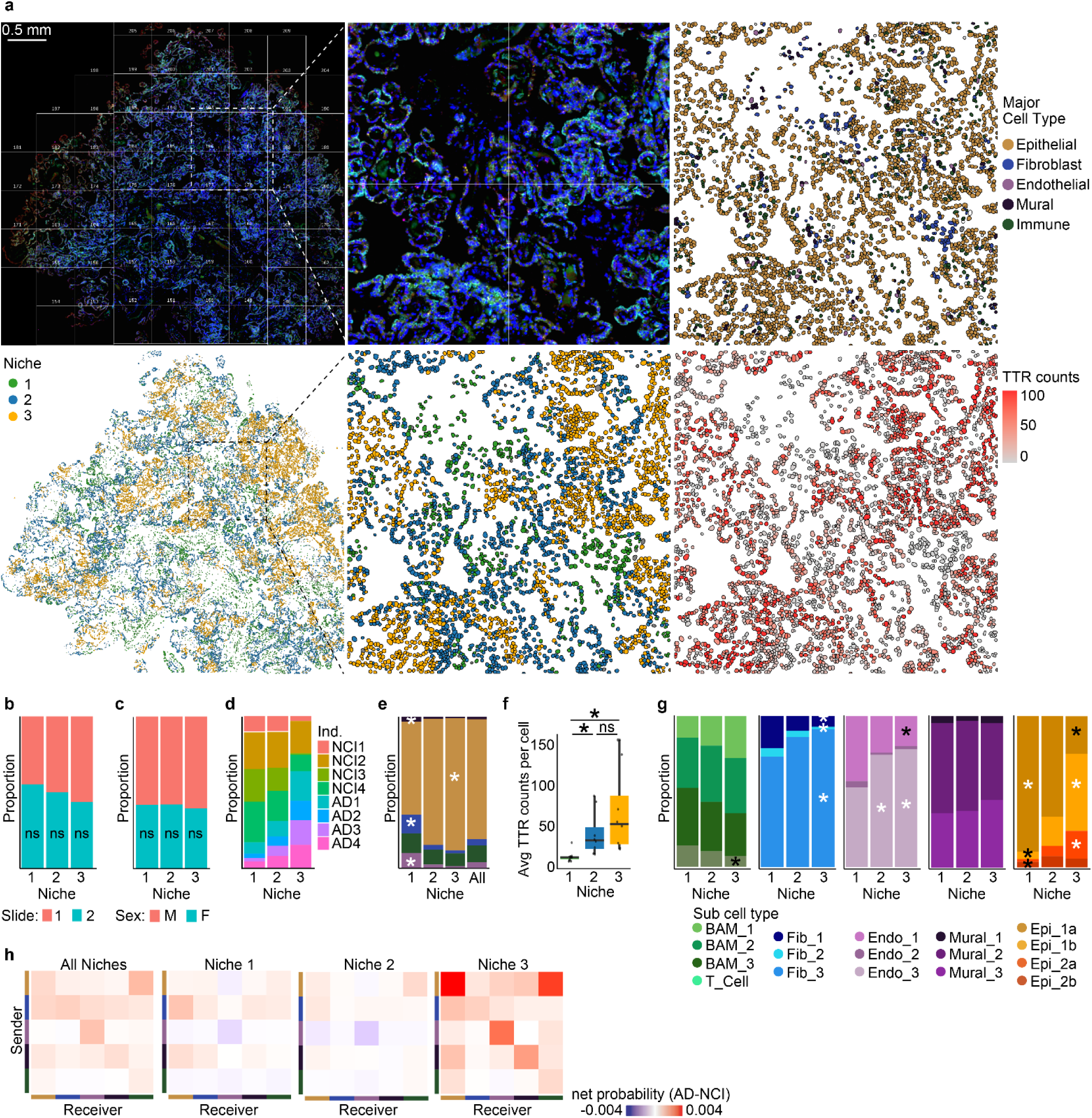
Niche-specific communication changes in Alzheimer’s disease. **a,** Example tissue section immunofluorescent stain (upper left) colored by niche (lower left). Middle: Higher magnification of 4 FOVs. Upper right: Major cell type annotation. Lower left: TTR counts per cell type. **b-e,** Stacked bar charts showing proportion of slide (b), sex (c), individual (d), major cell type (e) per niche. Linear mixed model on proportions (Mural, Fibroblast and Endothelial in niche 1, Epithelial in niche 3 * FDR < 0.05, ns: not significant). **f,** Boxplot shows individual- and niche-specific pseudobulk expression of TTR (* p <0.001, ns: not significant). **g,** Stacked bar charts showing subtype proportions per niche (* FDR < 0.05). **h,** AD differential CellChat net probability (CellChat net$prob) heatmaps among all ligand-receptor pairs in each colored sender-receiver pair, for all cells (left), or subset by niche.

## Extended Data Tables

**Extended Data Table 1**. Patient metadata

**Extended Data Table 2**. SN major celltype limma and GO

**Extended Data Table 3**. SN major and sub proportions and limma

**Extended Data Table 4**. ST major celltype limma and GO

**Extended Data Table 5**. ST major and sub proportions and limma

**Extended Data Table 6**. ATAC major celltype limma and GO

**Extended Data Table 7**. Epi SN subcelltype limma and GO

**Extended Data Table 8**. Epi ST subcelltype limma and GO

**Extended Data Table 9**. Epi ATAC subcelltype limma and GO

**Extended Data Table 10**. Epi SN major and sub cogdx and pattern limma and GO

**Extended Data Table 11**. Epi SN major and sub cog_dec limma and GO

**Extended Data Table 12**. Epi ST major cogdx limma and GO

**Extended Data Table 13**. Epi ATAC TF motifs subtype limma

**Extended Data Table 14**. CSF proteomic DEG limma and GO

**Extended Data Table 15**. Imm SN subcelltype limma and GO

**Extended Data Table 16**. Imm ATAC subcelltype limma and GO

**Extended Data Table 17**. Imm SN major and sub cogdx and pattern limma and GO

**Extended Data Table 18**. Imm ST major and sub cogdx

**Extended Data Table 19**. Fib SN subcelltype limma and GO

**Extended Data Table 20**. Fib SN major and sub cogdx and pattern limma and GO

**Extended Data Table 21**. Fib ST major cogdx limma and GO

**Extended Data Table 22**. Endo SN subcelltype limma and GO

**Extended Data Table 23**. Mural SN subcelltype limma and GO

**Extended Data Table 24**. Endo SN major cogdx limma and GO

**Extended Data Table 25**. Endo ST major cogdx limma and GO

**Extended Data Table 26**. Mural SN major cogdx limma and GO

**Extended Data Table 27**. Mural ST major cogdx limma and GO

**Extended Data Table 28**. Proteomic DEG limma and GO

**Extended Data Table 29**. SN and ST CellChat net_pathways and limma results

**Extended Data Table 30**. Canonical markers and references

## Notes

### Competing Interest Statement

The authors have declared no competing interest.

### Summary of Updates

Version 2 includes updated figures correcting display and alignment issues introduced during figure assembly in Illustrator. These changes do not affect the underlying data, analyses, or conclusions.

